# APOBEC3A drives acquired resistance to targeted therapies in non-small cell lung cancer

**DOI:** 10.1101/2021.01.20.426852

**Authors:** Hideko Isozaki, Ammal Abbasi, Naveed Nikpour, Adam Langenbucher, Wenjia Su, Marcello Stanzione, Heidie Frisco Cabanos, Faria M. Siddiqui, Nicole Phan, Pégah Jalili, Sunwoo Oh, Daria Timonina, Samantha Bilton, Maria Gomez-Caraballo, Hannah L. Archibald, Varuna Nangia, Kristin Dionne, Amanda Riley, Matthew Lawlor, Mandeep Kaur Banwait, Rosemary G. Cobb, Lee Zou, Nicholas J. Dyson, Christopher J. Ott, Cyril Benes, Gad Getz, Chang S. Chan, Alice T. Shaw, Jessica J. Lin, Lecia V. Sequist, Zofia Piotrowska, Jeffrey A. Engelman, Jake June-Koo Lee, Yosef Maruvka, Rémi Buisson, Michael S. Lawrence, Aaron N. Hata

## Abstract

Acquired drug resistance to even the most effective anti-cancer targeted therapies remains an unsolved clinical problem. Although many drivers of acquired drug resistance have been identified^1‒6^, the underlying molecular mechanisms shaping tumor evolution during treatment are incompletely understood. The extent to which therapy actively drives tumor evolution by promoting mutagenic processes^7^ or simply provides the selective pressure necessary for the outgrowth of drug-resistant clones^8^ remains an open question. Here, we report that lung cancer targeted therapies commonly used in the clinic induce the expression of cytidine deaminase APOBEC3A (A3A), leading to sustained mutagenesis in drug-tolerant cancer cells persisting during therapy. Induction of A3A facilitated the formation of double-strand DNA breaks (DSBs) in cycling drug-treated cells, and fully resistant clones that evolved from drug-tolerant intermediates exhibited an elevated burden of chromosomal aberrations such as copy number alterations and structural variations. Preventing therapy-induced A3A mutagenesis either by gene deletion or RNAi-mediated suppression delayed the emergence of drug resistance. Finally, we observed accumulation of A3A mutations in lung cancer patients who developed drug resistance after treatment with sequential targeted therapies. These data suggest that induction of A3A mutagenesis in response to targeted therapy treatment may facilitate the development of acquired resistance in non-small-cell lung cancer. Thus, suppressing expression or enzymatic activity of A3A may represent a potential therapeutic strategy to prevent or delay acquired resistance to lung cancer targeted therapy.

---

Large-scale genomic analyses of non-small-cell lung cancer (NSCLC) cohorts have defined the landscape of mutational processes that contribute to tumor development^9^, revealing extensive inter- and intra-tumoral heterogeneity^10^ that in principle may lead to incomplete therapy response and drug resistance^11^. By comparison, less is known about mutational mechanisms that are operative in lung cancers that respond to targeted therapies and subsequently relapse due to the development of acquired drug resistance. While pre-existing resistant clones may emerge under the selective pressure of therapy, cancer cells may also adapt and evolve in response to treatment. Epigenetic changes can facilitate drug tolerance and tumor cell survival during therapy^12,13^, with subsequent evolution driven by *de novo* acquisition of genomic resistance mechanisms in some cases^14,15^. This potential for ongoing evolution of resistant clones during therapy is demonstrated by NSCLC patients whose tumors accumulated compound mutations over the course of treatment with sequential tyrosine kinase inhibitor (TKI) targeted therapies^16,17^. In this study, we investigated whether specific mutational mechanisms drive genomic evolution of lung cancers during treatment with targeted therapies.

## Evolving EGFR-mutant NSCLC cells accumulate APOBEC mutations

To identify mutational signatures that reflect tumor evolution during therapy, we compared resistant EGFR-mutant NSCLC clones that acquired the *EGFR^T790M^* gatekeeper resistance mutation via evolution from persistent drug-tolerant tumor cells during gefitinib treatment to those derived from pre-existing *EGFR^T790M^* cells^14^ (Figure 1a). Whole-genome sequencing revealed that resistant clones that evolved from persistent drug-tolerant cells (“late”-evolving) harbored more single-nucleotide variants (SNVs) compared to the parental cell population than “early” pre-existing resistant clones (Figure 1b). We resolved mutational signatures within each clone using non-negative matrix factorization (NMF). Late clones harbored significantly more APOBEC cytidine deaminase mutations (Sanger Signature 2/13) (Figure 1c-d, Extended Data Figure 1a). Consistent with the NMF analysis, late clones harbored significantly more T<u>C</u>T→T<u>G</u>T and T<u>C</u>A→T<u>G</u>A mutations that are highly specific for APOBEC^18^ (Extended Data Figure 1b). Additionally, we observed increased kataegis^19^ (mutation clusters) in late clones, a hallmark of APOBEC mutagenesis (Figure 1e). Next, we reconstructed the evolutionary history of resistant clones. The majority (>70%) of new mutations in early clones were shared with other early clones, defining a recent common ancestor that had acquired *EGFR^T790M^* prior to treatment (Extended Data Figure 1c-d). In contrast, the late resistant clones harbored almost exclusively (>99%) new private mutations, consistent with independent clonal evolution both before and during TKI treatment. NMF analysis revealed increased APOBEC signatures in the private mutations in late clones, compared to the shared or private mutations in early clones (Extended Data Figure 1e), suggesting that APOBEC mutagenesis coincided with the period of TKI treatment (Extended Data Figure 1f). To test this hypothesis, we derived two PC9 single-cell clones and treated them with gefitinib until they evolved *EGFR^T790M^*. Whole-genome sequencing of the matched pre-treatment and resistant clones revealed an increased number of APOBEC mutations and associated kataegis events after TKI treatment (Figure 1f).

**Figure 1.**
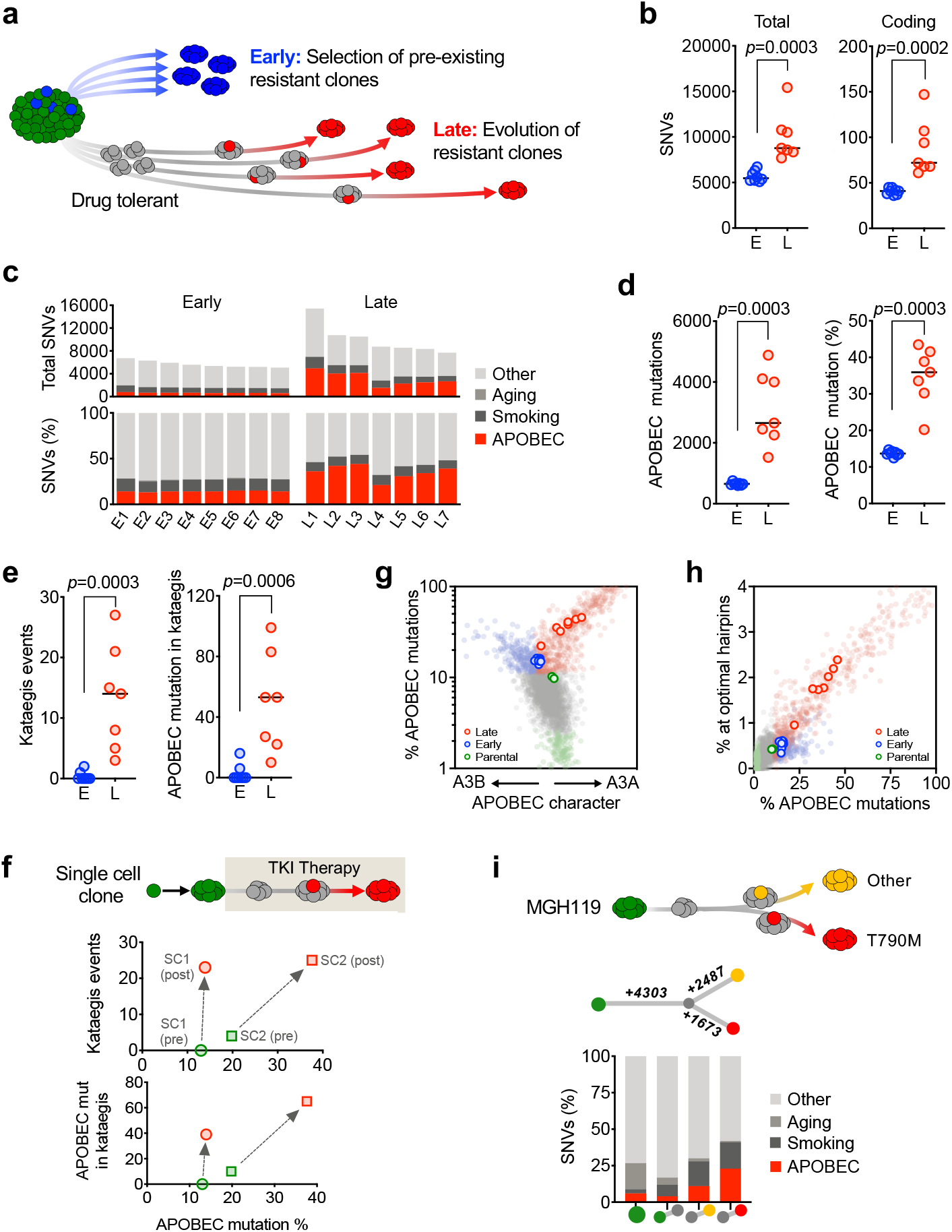
TKI-resistant clones that evolve from drug-tolerant persister cells accumulate APOBEC mutations. **a**, Evolutionary pathway of EGFR^T790M^ resistant PC9 clones. **b**, Number of new single-nucleotide variants (SNVs) in PC9 early and late resistant clones compared to parental cells. **c**, New SNVs attributed to mutational signatures in early and late resistant clones. **d**, APOBEC signature SNVs in early and late resistant clones. **e**, Kataegis mutation clusters and associated APOBEC mutations in early and late resistant clones. **f**, Pre- and acquired post-treatment kataegis mutation clusters and associated APOBEC mutations in matched treatment naive (green) and TKI-resistant (red) PC9 single-cell clones. Acquired post-treatment mutations were defined as private mutations only present in the resistant clone. **g**, A3A (YTC) and A3B (RTC) character of parental (green open circles), early (blue open circles) and late (red open circles) resistant clones. A reference set of ~2600 WGS-analyzed tumors from the International Cancer Genome Consortium (ICGC) Pan-Cancer Analysis of Whole Genomes (PCAWG) project is plotted for comparison. Samples were designed APOBEC+ if ≥10% of total mutations were assigned to the APOBEC mutational signature by NMF analysis, then classified according to A3A vs. A3B character: A3A-dominated samples (enrichment of mutations at YTC trinucleotides) are colored red, A3B-dominated samples (enrichment of mutations at RTC trinucleotides) are colored blue. APOBEC-samples (<1% APOBEC mutations) are colored green. Samples with 1-10% APOBEC mutations were colored grey. **h**, Percentage of APOBEC mutations in parental, early and late resistant clones that map to optimal A3A hairpin motifs. PCAWG reference samples are colored as in panel g. **i**, Evolutionary pathways and associated APOBEC mutational signatures of MGH119 TKI-resistant clones.

Several APOBEC family members have been implicated in tumor evolution, most prominently APOBEC3A (A3A) and APOBEC3B (A3B)^20^. Recent studies have demonstrated context-specific preferences of A3A and A3B at TpC sites: A3A exhibits preference for a pyrimidine at the −2 position (YTC) whereas 3B prefers a purine (RTC)^21^. We calculated the YTC vs RTC character of mutations in the early and late clones and compared these to a cohort of ~2600 WGS-analyzed tumors from the International Cancer Genome Consortium (ICGC) Pan-Cancer Analysis of Whole Genomes (PCAWG) project^22^ (Figure 1g). Whereas early clones showed mid-range levels of APOBEC mutations (~10% of total mutations), without clear dominance of either A3A or A3B, late clones exhibited elevated APOBEC content (~40%) with a striking A3A (YTC) bias. To provide further evidence supporting a role for A3A, we examined whether APOBEC mutations occurred in the context of hairpin loop secondary structures that were recently shown to be highly specific for A3A^23^. Similar to A3A-positive tumors from PCAWG, late clones harbored significantly increased numbers of mutations in optimal hairpins (Figure 1h), further illustrating the role of A3A in the evolution of these PC9 clones.

To extend these findings, we examined an independent model of acquired EGFR TKI resistance. We treated MGH119 cells, which do not harbor pre-existing *EGFR^T790M^* cells^14^, with gefitinib until two independent resistant clones emerged, one of which harbored *EGFR^T790M^*. Compared to the parental cell line, ~4300 unique mutations were present in both resistant clones, suggesting they evolved from a common ancestor. In addition, each clone harbored an additional 2487 and 1673 private mutations, respectively (Figure 1i). Mutational signature analysis revealed an elevated frequency of APOBEC mutations in the private branches, consistent with accumulation of APOBEC mutations during TKI treatment. Similar to the PC9 clones, the APOBEC mutations in MGH119 resistant cells exhibited A3A character (Extended Data Figure 1g). Together, these results reveal that accumulation of A3A mutations commonly occurs in evolutionary models of EGFR TKI acquired resistance.

## Lung cancer targeted therapies induce APOBEC3A mutagenesis

To establish a causal relationship between TKI therapy and APOBEC mutagenesis, we examined whether TKI treatment induces expression of APOBEC family genes. Upon gefitinib treatment, we consistently observed rapid up-regulation of A3A, but not other APOBEC cytidine deaminases, that was sustained in drug-tolerant persister cells after 14 days of treatment (Figure 2a). Analysis of an independent RNA-seq dataset also revealed increased expression of A3A but not other APOBEC family members after EGFR TKI treatment (Extended Data Figure 2a). Similar results were observed upon treatment of cells with the third-generation EGFR TKI osimertinib, which has recently become the standard first-line therapy for EGFR-mutant NSCLC^24^ (Extended Data Figure 2b). We extended these findings to an expanded panel of 13 EGFR NSCLC cell lines including patient-derived cell lines generated from pre-treatment, on-treatment or progression biopsies (with acquired *EGFR^T790M^*). Altogether, we observed >4-fold induction of A3A expression after osimertinib treatment in 10/13 models representing all three clinical scenarios (Figure 2b), whereas A3B expression was relatively unchanged. To gain insight into the mechanism of TKI-induced A3A expression, we performed ATAC-seq on PC9 cells after 2 weeks of TKI treatment and observed three regions of increased chromatin accessibility upstream of the A3A transcriptional start site (Extended Data Figure 2c). Examination of the ENCODE database identified several transcription factors that have been shown to bind at these putative enhancer regions including NF-kB, c-Jun/Fos/AP-1, STAT2/STAT3 and the ETS transcription factor SPI-1. Consistent with prior reports demonstrating NF-kB activation in the response to EGFR TKI treatment^25,26^, we observed a global increase in chromatin accessibility at NF-kB motifs, but not sites bound by the other transcription factors (Extended Data Figure 2d). Knockdown of components of the NFKB signaling cascade reduced TKI-induced APOBEC expression (Extended Data Figure 2e). Thus, induction of A3A expression is a common response of EGFR-mutant NSCLC cells to TKI treatment, likely a consequence of activation of innate immune signaling.

**Figure 2.**
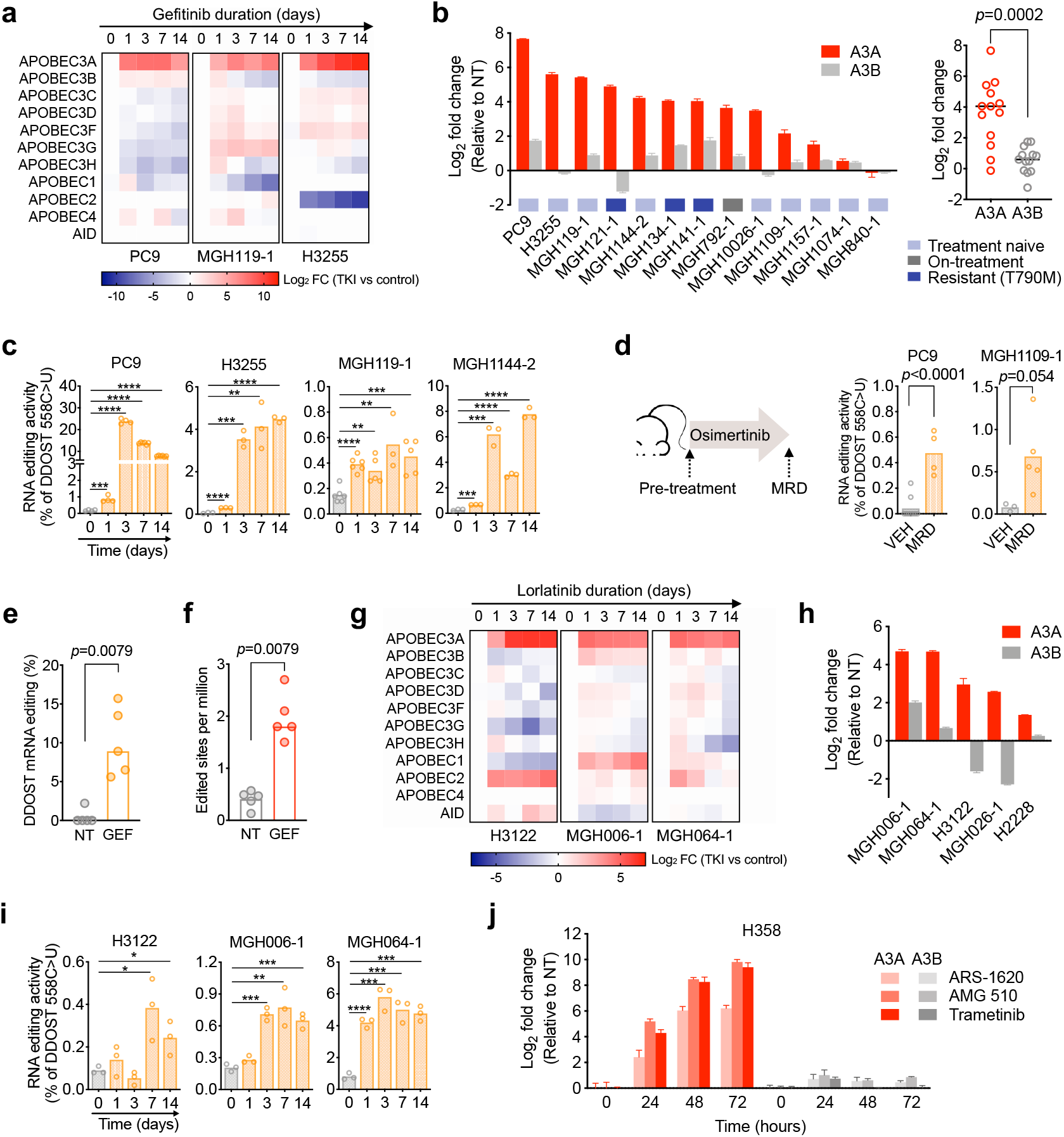
Lung cancer targeted therapies induced APOBEC3A mutagenesis. **a**, EGFR-mutant NSCLC cell lines were treated with 1 μM gefitinib for up to 14 days and gene expression was determined by quantitative RT-PCR (3 biological replicates). **b**, Expression of A3A and A3B in EGFR-mutant NSCLC patient-derived cell lines after treatment with 1 μM osimertinib for 24 hours (3 biological replicates). **c**, DDOST mRNA editing in EGFR-mutant NSCLC cell lines treated with 1 μM osimertinib for up to 14 days (3 biological replicates). **d**, DDOST mRNA editing in TKI-treated EGFR NSCLC xenograft tumors. Mice were treated with osimertinib until tumors had regressed to a stable minimal residual disease (MRD) state. **e**, Edited DDOST mRNA-seq reads from PC9 cells treated with 300 nM gefitinib for 14 days. **f**, Transcriptome-wide editing of A3A hairpin motifs in mRNA-seq reads from PC9 cells treated with 300 nM gefitinib for 14 days. **g**, Expression of APOBEC family genes in ALK fusion-positive NSCLC cells after treatment with 100 nM lorlatinib as determined by quantitative RT-PCR (3 biological replicates). **h**, Expression of A3A and A3B in ALK fusion-positive NSCLC cell lines treated with 100 nM lorlatinib for 24 hours (3 biological replicates). **i**, DDOST mRNA editing in ALK fusion-positive NSCLC cell lines treated with 100 nM lorlatinib for up to 14 days (3 biological replicates). **j**, Expression of A3A and A3B in H358 *KRASG12C* NSCLC cells treated with 1 μM AMG 510, 1 μM ARS-1620 or 100 nM trametinib for up to 72 hours. Data are normalized to untreated control (3 biological replicates). **c, i**Two-tailed Student’s *t*-test; \*\**p* < 0.01, \*\*\**p* < 0.001, \*\*\*\**p* < 0.0001.

Next, we investigated whether TKI-induced A3A expression is accompanied by increased mutagenic activity. To specifically assess A3A activity, we employed two complementary approaches that take advantage of the tendency of A3A to edit RNA and DNA at TpC dinucleotides presented within defined stem-loop hairpin motifs^23,27^. Importantly, these methods enable discrimination of A3A from A3B editing activity^27^. First, we quantified A3A C>T editing at a hairpin motif within the DDOST gene using an allele-specific droplet digital PCR assay ^27^ (Extended Data Figure 3a-c). Consistent with the induction of A3A expression, DDOST editing was significantly increased in EGFR-mutant NSCLC cell lines shortly after initiation of TKI treatment and remained elevated in drug-tolerant persister cells after 14 days of treatment (Figure 2c). To corroborate these *in vitro* findings, we treated mice bearing PC9 or EGFR-mutant NSCLC patient-derived xenograft (PDX) tumors with osimertinib until tumors regressed to a stable minimal-residual-disease state (Extended Data Figure 3d). DDOST RNA editing was increased in TKI-treated residual tumors compared with untreated tumors (Figure 2d), confirming that TKI treatment induces A3A editing in persistent tumor cells *in vivo*. As a second approach, we examined whether A3A editing could be detected by RNA-seq by quantifying C>U mutations in transcripts corresponding to A3A hotspots^27^. Both TKI treatment as well as overexpression increased C>U edited DDOST transcripts in PC9 cells (Figure 2e, Extended Data Figure 3e). Additionally, we developed a computational tool called ApoTrack that integrates reads containing UCN > UUN mutations in hairpin loops of sequence NUC or NNUC at the end of stably paired stems at ~2000 sites across the transcriptome. This revealed widespread APOBEC3A RNA editing in TKI treated cells (Figure 2f, Extended Data Figure 3f).

**Figure 3.**
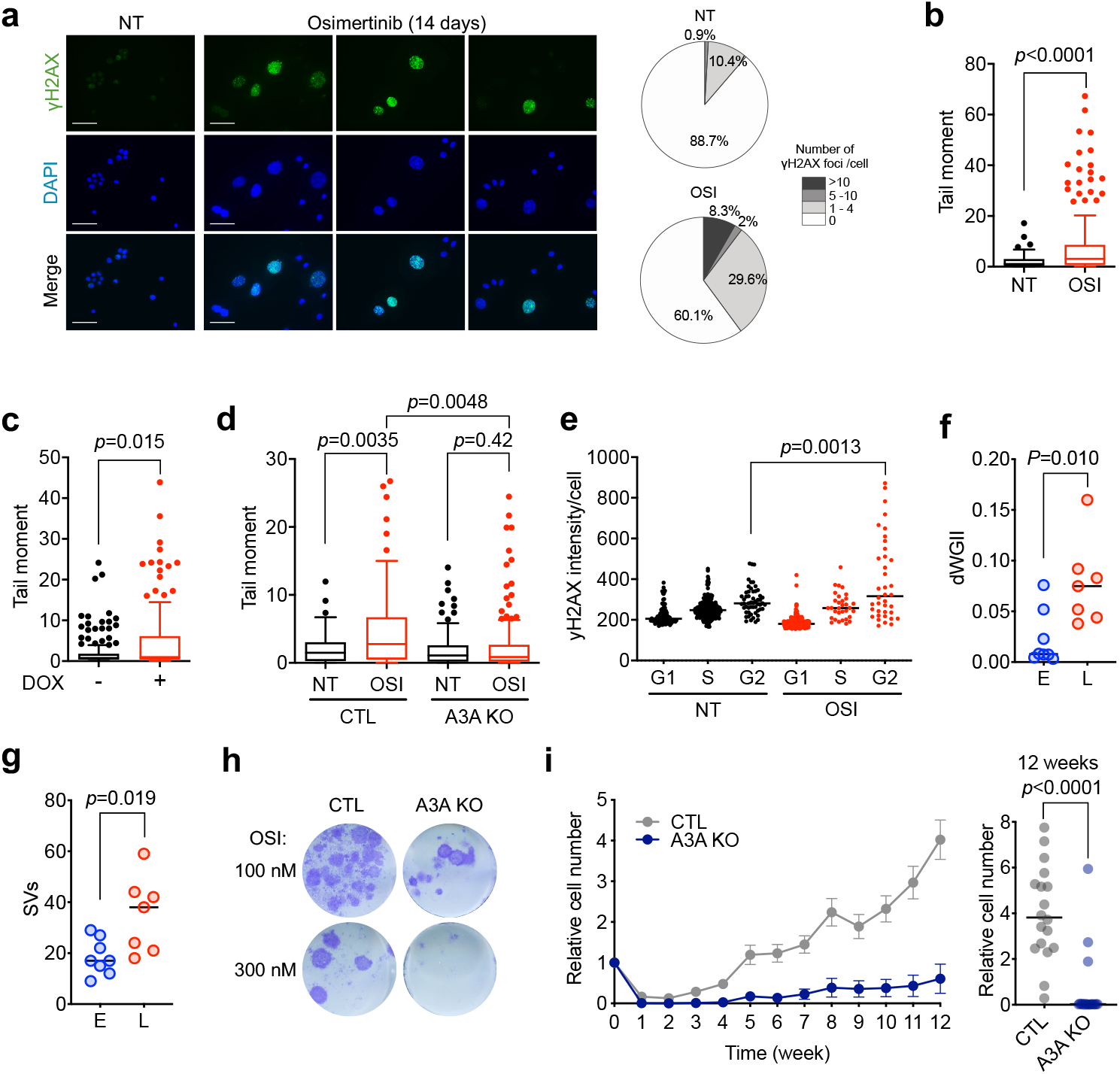
TKI-induced APOBEC3A leads to genomic instability and facilitates evolution of drug-resistant clones. **a**, Representative immunofluorescence images of γH2AX in PC9 cells treated with or without 1 μM osimertinib for 14 days. NT, no treatment; scale bars = 70 μm. Pie charts show percentage of cells containing γH2AX foci. **b**, PC9 cells were treated with or without osimertinib (OSI) for 2 weeks and DSBs were assessed by neutral comet assay. Data are represented as Tukey box and whisker plots and are representative of three independent experiments. **c**, PC9 cells transduced with a doxycycline-inducible wild-type A3A were treated with doxycycline to induce expression of A3A for 96 hours prior to being analyzed by comet assay. Data are representative of three independent experiments. **d**, PC9 control and PC9 A3A knockout (KO) cells were treated with or without 1 μM osimertinib for 14 days and DSBs were assessed by neutral comet assay. Data are representative of three independent experiments. **e**, PC9 cells were treated with 1 μM osimertinib for 0 or 14 days and stained for γH2AX to quantify DNA damage, and EdU/DAPI to resolve cell cycle phase (see Extended Data Figure 5c). Data shown are representative of two independent experiments. **f**, Late-evolving PC9 resistant clones have increased copy number changes (as quantified by differential Weighted Genomic Integrity Index; dWGII) compared to early clones. **g**, Late-evolving clones have increased number of structural variations. **h**, PC9 A3A KO and control cells were treated with osimertinib for 6 weeks and colonies of drug-tolerant persister cells were visualized by crystal violet. **i**, PC9 A3A KO and control cells were treated with 100 nM osimertinib for 12 weeks and cell number was monitored using RealTime-Go (N=18 independent pools each, mean ± s.e.m.). Right panel shows the cell number of each pool at week 12.

To place these findings into the evolutionary context of acquired drug resistance (Figure 1a), we examined A3A expression and editing activity during the evolutionary trajectories of resistant *EGFR^T790M^* PC9-GR2 (early) and PC9-GR3 (late) cells that we previously described ^14^. Despite having previously accumulated APOBEC mutations (Figure 1c), the baseline A3A expression and activity of PC9-GR3 cells were similar to untreated parental PC9 cells and early-evolving PC9-GR2 cells (Extended Data Figure 4a-b), indicating that the A3A mutagenesis induced by TKI during the evolutionary pathway had resolved. Recent studies of lung cancer patient cohorts as well as *in vitro* cultured tumor cells have revealed that APOBEC editing can occur sporadically, likely in response to episodic stimuli^28-30^. We hypothesized that suppression of oncogenic signaling by TKI might provide a transient stimulus in persistent drug-tolerant clones that is ultimately resolved if oncogenic signaling is restored, for instance with acquisition of a secondary EGFR resistance mutation such as *EGFR^T790M^*. Consistent with this notion, when TKI-treated persistent drug-tolerant cells were removed from drug, A3A expression returned to baseline (Extended Data Figure 4a-b). Furthermore, suppression of EGFR^T790M^ with a third-generation EGFR TKI for 24 hours re-induced A3A expression and editing activity in PC9-GR3 cells (Extended Data Figure 4a). Importantly, A3A expression and editing was also induced in PC9-GR2 upon treatment with a third-generation TKI, indicating that the lack of APOBEC mutations in this resistant clone was due not to an intrinsic inability to induce A3A, but rather to the lack of an effective stimulus. Thus, A3A expression and activity is tightly linked with suppression of oncogenic EGFR signaling throughout the evolutionary trajectory of EGFR-mutant NSCLC clones (Extended Data Figure 4c).

**Figure 4.**
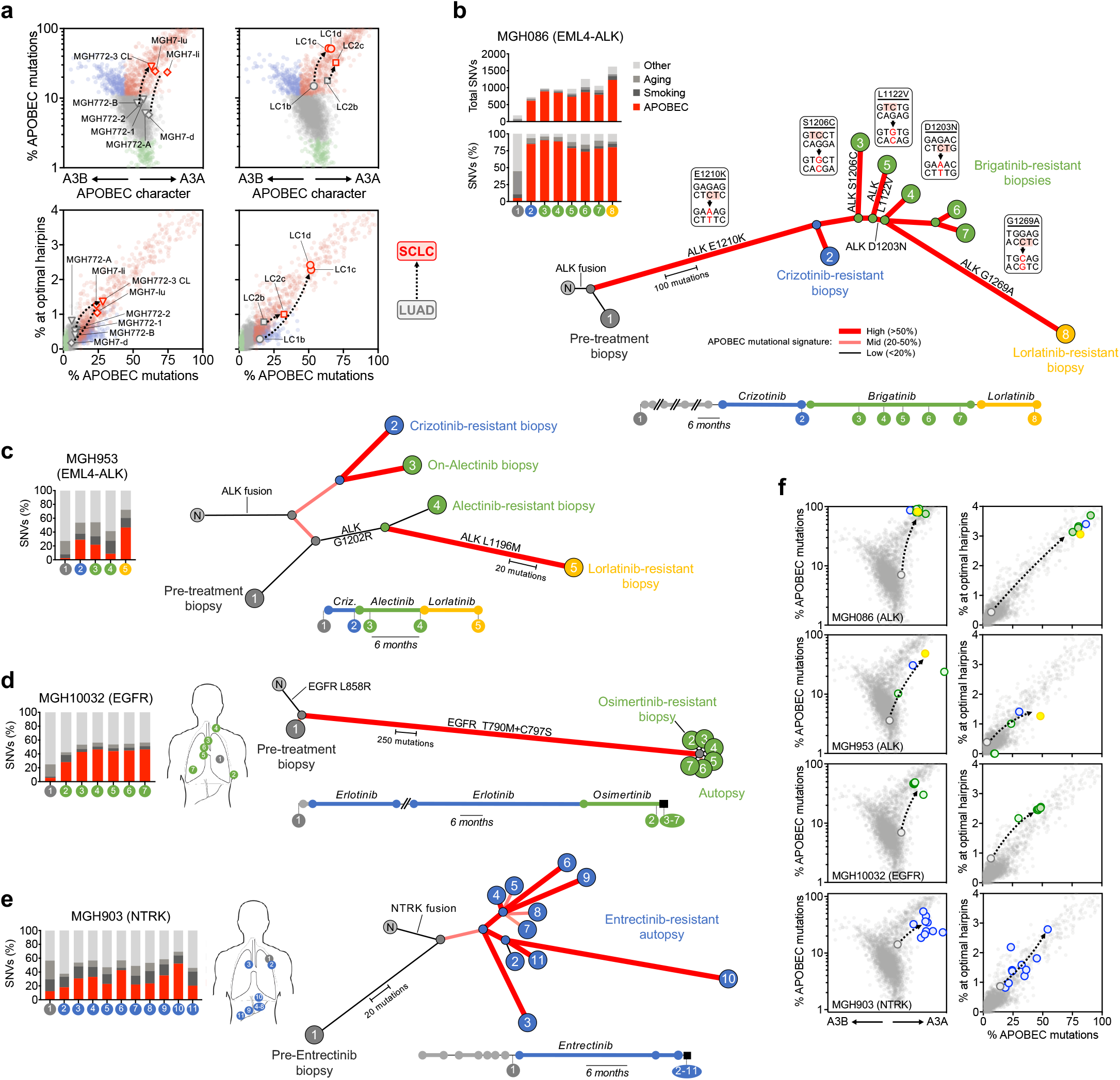
APOBEC3A mutational signatures in NSCLC patients with acquired TKI resistance. **a**, A3A (YTC) and A3B (RTC) character (upper panels) and A3A hairpin motif mutations (lower panel) of SCLC-transformed (red symbols) and corresponding adenocarcinoma NSCLCs (gray symbols). Similar to Figure 1g-h, ~2600 WGS-analyzed tumors from the PCAWG project are plotted for comparison. Tumors from the same patient are connected by a dashed line. Clinical histories for MGH772 sequential biopsies and MGH7 autopsy shown in the left panel are depicted in Extended Data Figure 8a-b. Right panels depict pre- and post-transformation tumors described in Lee et al.^40^ CL = patient-derived cell line, lu = lung metastasis, li - liver metastasis, d = diaphragm metastasis. **b**, Mutational signatures and clonal relationship of tumor biopsies from patient MGH086 over the course of sequential ALK TKI therapy17. Branches depict the fraction of mutations corresponding to an APOBEC signature. Base level mutations are indicated for each resistance mutation with TpC APOBEC motif highlighted. **c**, Mutational signatures and clonal relationship of multiple sequential biopsies from patient MGH953 over the course of sequential ALK TKI therapy^17^. **d**, Mutational signatures and clonal relationship of pre-/post-treatment biopsies and autopsy sites from patient MGH10032 after sequential EGFR TKI therapy. **e**, Mutational signatures and clonal relationship of pre-entrectinib biopsy and autopsy samples from patient MGH903. **f**, A3A (YTC) and A3B (RTC) character (left panels) and A3A hairpin motif mutations (right panels) of clinical cases shown in b-e. Samples are colored according to sequential TKI treatment history, and dotted lines indicate clinical trajectories. Gray cloud indicates WGS-analyzed tumors from the PCAWG project.

Given the similarities between signaling pathways activated by different driver oncogenes in lung cancer, we investigated whether other lung cancer targeted therapies similarly induce A3A activity. Treatment of ALK fusion-positive lung cancer cell lines with the third-generation ALK inhibitor lorlatinib induced A3A expression (Figure 2g-h) and RNA editing activity (Figure 2i). Similarly, treatment of KRAS-mutant NSCLC cell lines with the KRAS^G12C^ inhibitors ARS-1620 and AMG 510 or the MEK inhibitor trametinib induced A3A expression (Figure 2j) Finally, we performed RNA-seq on H358 cells treated with ARS-1620 or trametinib and observed an increase in A3A editing at the DDOST hairpin hotspot as well as across the transcriptome (Extended Data Figure 3g-h). Taken altogether, these results reveal that induction of A3A expression and activity is a common response of oncogene-driven lung cancer cells treated with targeted therapies

## TKI-induced APOBEC3A causes DNA damage and drives evolution of drug resistance

Prior studies have demonstrated that A3A mutagenesis can cause double-strand DNA breaks (DSBs) and activate DNA damage response (DDR) signaling^31–33^. We observed elevated Ser139 phosphorylation of the histone variant H2AX, a marker of the DNA damage response^34^, in PC9 cells after 2 weeks of osimertinib treatment (Figure 3a). Consistent with this, neutral comet assay revealed increased DSBs in these cells (Figure 3b). To investigate whether A3A contributes to DNA damage sustained during TKI treatment, we first confirmed that overexpression of A3A is sufficient to cause DSBs using a doxycycline-inducible A3A overexpression construct (Figure 3c, Extended Data Figure 3b-c). We next assessed whether A3A expression is necessary for TKI-induced DSBs. After 14 days of osimertinib treatment, DSBs were significantly reduced in PC9 cells with CRISPR-mediated deletion of A3A (A3A KO) compared with control cells (Figure 3d, Extended Data Figure 5a). Prior studies have revealed that APOBEC mutational signatures exhibit lagging-strand bias reflecting APOBEC mutagenesis of single-strand DNA during replication^35,36^. Consistent with these findings, APOBEC mutations in late-evolving resistant PC9 clones exhibited lagging-strand asymmetry (Extended Data Figure 5b), indicating that APOBEC editing of DNA had occurred in replicating cells. Although TKI-treatment induces G1 arrest, over time a subset of drug-tolerant cells resume cell division and begin to proliferate^12^. Mapping γH2AX onto the cell cycle distribution of TKI-treated PC9 cells revealed that γH2AX was most prominently localized to a subpopulation of cells that had resumed cell division and were in G2 phase (Figure 3e, Extended Data Figure 5). Thus, TKI treatment induces A3A-catalyzed genomic damage in proliferating drug-tolerant persister cells.

Error-prone repair of double-strand DNA breaks or mitotic errors resulting from replicating cells containing un-repaired breaks can increase chromosomal instability, leading to genomic heterogeneity that facilitates tumor evolution^37^. To determine the genomic sequelae of A3A-induced DSBs in emerging drug-resistant clones, we compared chromosomal aberrations in early- and late-evolving PC9 resistant clones. First we identified copy-number changes in resistant clones relative to the parental cells, calculating a differential version of the Weighted Genomic Integrity Index (WGII)^38^, which we call dWGII. This revealed significantly increased copy number changes in late-evolving clones, with fewer changes in early-evolving clones (Figure 3f, Extended Data Figure 6a-b). Second, we quantified structural variations (SVs), including translocations, inversions, and large insertions/deletions, using dRanger and Breakpointer^39^, again relative to the parental cells. This demonstrated increased SVs in late-evolving clones and fewer SVs in early-evolving clones (Figure 3g, Extended Data Figure 6c). These data suggest that lung cancer targeted therapies can increase the genomic damage that leads to chromosomal instability in evolving drug resistant cells (Extended Data Figure 6d).

Finally, we examined whether A3A-induced DNA damage facilitates the evolution of acquired resistance to targeted therapy. PC9 A3A KO or control cells were treated with osimertinib and monitored for the emergence of drug-tolerant and resistant clones. Deletion of A3A suppressed the emergence of drug-tolerant colonies (Figure 3h) and subsequent evolution of drug-resistant clones over long-term TKI treatment (Figure 3i). To rule out the possibility of off-target genomic effects of CRISPR, we independently confirmed that suppression of A3A using shRNA also suppressed the evolution of drug-resistant clones (Extended Data Figure 7). Taken all together, these results indicate that therapy-induced A3A can drive genomic instability in persistent drug-tolerant cells and facilitate the evolution of resistant clones that harbor increasing genomic heterogeneity of point mutations and structural variations.

## TKI-resistant patients harbor APOBEC mutational signatures

Recently, it has been reported that transformation of EGFR-mutant lung cancers from adenocarcinoma to small-cell lung cancer (SCLC) histology at the time of acquired resistance is associated with the appearance of APOBEC mutational signatures^40,41^. Similarly, we detected APOBEC mutational signatures in two previously reported cases of SCLC transformation from our group^42,43^ (Extended Data Figure 8a-b). These APOBEC mutational signatures were observed in the SCLC-transformed tumors but not in the corresponding patients’ non-transformed adenocarcinomas. Further analysis confirmed that the APOBEC mutations in each of these transformed tumors, as well two cases reported by Lee et al.^40,41^, exhibited clear A3A character, consistent with our experimental models of TKI-induced A3A mutagenesis (Figure 4a).

As our pre-clinical data suggested that induction of A3A mutagenesis is associated with the evolutionary pathway rather than the specific mechanism of acquired resistance, we hypothesized that A3A mutagenesis in SCLC-transformed tumors might reflect persister evolution, in contrast to other resistance mechanisms such as *EGFR^T790M^*, which has been detected at low levels in some patients at the time of diagnosis and may emerge from pre-existing resistant clones^44,45^. To investigate whether APOBEC mutations accumulate in tumors that evolve mechanisms of resistance during treatment, we examined a trio of ALK fusion-positive NSCLC cases that acquired compound ALK resistance mutations after treatment with multiple ALK inhibitors^17^ (Extended Data Figure 8c, 9a-b). Importantly, the stepwise development of compound mutations indicates continued linear evolution of the same resistant clone in response to sequential TKI therapy. In two of the three patients, we observed increased APOBEC mutational signatures in resistant tumors after TKI treatment (Figure 4b-c, Extended Data Figure 8d). In one case (MGH086), each of the ALK resistance mutations resulted from C->T or C->G substitutions at TpC motifs, suggesting that they resulted from on-going APOBEC mutagenesis during TKI therapy. Additionally, we performed whole-genome sequencing on multiple metastatic sites from a rapid autopsy of an EGFR-mutant NSCLC patient who developed compound *EGFR^T790M/C797S^* mutations after an exceptional response to sequential EGFR TKI therapy (Figure 4d, Extended Data Figure 8e, 9c). Each metastatic site exhibited a similar mutational profile dominated by a dramatic increase in APOBEC mutations, consistent with evolution of a single common *EGFR^T790M/C797S^* clone. By comparison, no increase in APOBEC mutational signatures was observed in metastatic sites from a patient with a shorter response to EGFR TKIs and little evidence of clonal evolution during therapy (Extended Data Figure 8e, f). Finally, we examined multiple metastatic autopsy sites from a patient with an NTRK fusion-positive lung cancer after treatment with the TRK inhibitor entrectinib (Extended Data Figure 8g, 9d). In the post-entrectinib samples, we observed increased APOBEC mutational signatures that were not present in the pre-entrectinib biopsy (Figure 4e). Importantly, the pattern of APOBEC mutations defined three distinct subclones that emerged during therapy, suggesting independent APOBEC-driven clonal evolution of multiple clones during acquired resistance. Finally, we examined whether the APOBEC mutations observed in resistant tumors exhibited characteristics of A3A. In each of these clinical cases, the acquired APOBEC mutational signatures exhibited clear YTC bias and enrichment in optimal hairpin motifs, consistent with on-going A3A mutagenesis during TKI therapy (Figure 4f).

In summary, we find that commonly used lung cancer targeted therapies induce the expression and mutagenic activity of A3A. Sustained A3A activity in persistent drug-tolerant cells that survive initial TKI treatment leads to the accumulation of mutations and DNA damage, and ultimately may result in chromosomal aberrations such as copy-number changes and structural variations. We observe that a subset of patients harbors increased APOBEC mutations after TKI treatment, and in some cases, these mutations may be drivers of acquired drug resistance. Our results suggest that the acquisition of APOBEC mutational signatures after TKI therapy may be indicative of the evolutionary path of the resistant clone and provide a novel mechanism by which targeted therapies might unintentionally increase the adaptive mutability of cancer cells during treatment. We speculate that the genomic consequences of A3A activity may cooperate with other mechanisms of drug tolerance such as epigenetic chromatin changes and EMT. Thus, preventing expression or enzymatic activity of A3A may represent a potential therapeutic strategy to prevent or delay acquired resistance to lung cancer targeted therapy.

## Methods

### Cell lines and cell culture

All cell lines were listed in Extended Data Table 1. PC9, H3255, H3122 and H358 cells were obtained from the MGH Center for Molecular Therapeutics. The identity of these cell lines were verified by STR analysis (Bio-synthesis, Inc.) at the time that these studies were performed. Patient-derived cell lines were established in our laboratory from core biopsy or pleural effusion samples as previously described^42,46^. All patients signed informed consent to participate in a Dana-Farber–Harvard Cancer Center Institutional Review Board–approved protocol giving permission for research to be performed on their samples. All cell lines were maintained in RPMI (Life Technologies) supplemented with 10% FBS, except for H3255, MGH119-1 and MGH006-1 cells, which were maintained in ACL4 medium (Life Technologies) with 10% FBS. Gefitinib-resistant PC9-GR2, PC9-GR3 were established by culturing parental cells in escalating concentrations of gefitinib (10 nM-1 μM), as tolerated, as previously described. Early, drug-tolerant, late PC9 resistant clones were established by culturing in 300 nM gefitinib until resistant, at which point they were maintained in 1 μM gefitinib, as previously described^14^. MGH119 resistant clones were established by culturing MGH119-1 parental cells in 1 μM gefitinib until resistant clones emerged. During generation of resistance, medium and drug were replaced twice per week. PC9 resistant clones were established as previously described and listed in Extended Data Table 2^14^. All experiments were performed in RPMI with 10% FBS. All cells were routinely tested and verified to be free of mycoplasma contamination.

### Antibodies and reagents

For western blotting and immunofluorescence, the following antibodies were used: Flag-M2 (1:1000, Sigma-Aldrich #F1084), Actin (1:1000, Cell Signaling #4970), γH2AX (1:1000, Millipore #05-636). For cell culture studies, gefitinib, osimertinib, Lorlatinib, trametinib, ARS1620 (all from Selleck) and AMG 510 (Chemgood) were dissolved in DMSO to a final concentration of 10 mmol/L and stored at −20 °C. Unless otherwise specified, 1 μM concentration was used for in vitro cell culture experiments.

### Whole-genome and whole-exome DNA sequencing

Whole-genome sequencing (WGS) and whole-exome sequencing (WXS) was performed on DNA extracted from cell lines, snap-frozen tumor specimens or FFPE-preserved tumor tissue. Library preparation and Illumina paired-end sequencing (2 × 150 bp) was performed by the Broad Institute Genomics Platform. Sequencing was analyzed using the published Broad Institute pipelines^22,47^. Reads were aligned to the human genome (build hg19) using bwa^48^, then processed through the Picard pipeline (http://broadinstitute.github.io/picard) to recalibrate base quality scores, sort reads, mark duplicate reads, and perform indel realignment. Somatic mutations were called using MuTect^49^, and polymorphisms and mutations in parental samples were called using MuTect in unpaired mode. Somatic mutations were filtered to remove variants observed in parental samples. Analysis of shared and private mutations, and construction of sample phylogenies, was performed as previously^9,17,50,51^. Briefly, for each patient or cell line, all samples’ somatic mutations were combined into a single list, and supporting reads for each mutation were tabulated in each sample. Mutations were grouped into patterns of shared presence or absence across samples, and by shared clonal fraction. “Shared early” mutations in the PC9 clones were identified by being detectable (>=2 reads, >=1% of total reads) in all of the early-emerging samples. Clustered mutations (kataegis) were identified in each sample as runs of consecutive mutations (at least 3) separated from each other by <50 kb. Mutational signature analysis was performed as previously^23,52,53^. Samples from our cohort were aggregated with data from The Cancer Gene Atlas (TCGA)^54^ and the International Cancer Genome Consortium (ICGC) Pan-Cancer Analysis of Whole Genomes project (PCAWG)^22^, and non-negative matrix factorization (NMF) was used to decompose mutation lists into seven mutational signatures corresponding to known mutation processes including APOBEC, Aging, and Smoking (Extended Data Figure 1a). The fraction of mutations due to each signature was calculated from the NMF weighting factors. The fraction of mutations assigned to the APOBEC signature is shown on the y-axis of Figure 1g and x-axis of Figure 1h. APOBEC3A (A3A) and APOBEC3B (A3B) mutation character were measured as described previously^21,23,27^, by calculating the fraction of cytosine mutations occurring in the contexts YTC or RTC, respectively. To obtain a directional metric of each patient’s overall A3A vs. A3B mutation character, we calculated fracYTC minus twice fracRTC. This metric is shown on the x-axis of Figure 1g. Mutations at DNA hairpins were identified as described previously^23^, and the fraction of mutations occurring at optimal hairpins (expected relative mutation rate of ≥4) was calculated. This metric is shown on the y-axis of Figure 1h. Mutation strand asymmetry was analyzed as described previously^35^, calculating for each sample the log2 ratio of the count of mutations at cytosines located on the replicative lagging-strand template, and the count on the leading-strand template. Genomic copy number was analyzed as described previously^50^, with read depth, normalized to total sample coverage and segmented using circular binary segmentation. Purity and ploidy estimates were achieved by iteratively searching for the set of parameters that minimizes the total distance between each profile’s copy-number segments and the nearest integer copy-number state using Matlab’s fminsearch function. Purity of cell-line samples was fixed at 1. Copy ratio was corrected for purity and ploidy to yield total absolute copy number (Extended Data Figure 6a, left). Acquired copy-number changes were computed by subtracting the copy number of the parental sample (Extended Data Figure 6a, right). Genomic segments that changed in copy number between each derived sample and its parent were identified and plotted in Extended Data Figure 6b, and the extent of these changes was quantified as a single metric dWGII (Figure 3f), a differential version of the established Weighted Genomic Integrity Index metric^38^ defined as the fraction of the genome that either gained or lost at least one DNA copy, relative to the parental sample. Somatic structural variants (SVs) were identified using dRanger and BreakPointer as described previously^39,50^. SVs of different structural classes were counted (Extended Data Figure 6c) and visualized alongside mutation and copy-number data with CIRCOS plots^55^ (Extended Figure 6d) created using the Circa software (http://omgenomics.com/circa).

### RNA-seq analysis of mRNA editing activity

RNA sequencing data was aligned to the hg19 genome and transcriptome using the STAR aligner^56^. A3A-mediated mRNA editing was monitored using the RNA hairpin hotspot in the gene *DDOST* as described previously^27^, calculating in each sample the percentage of reads carrying a G->A change at chr1:20981977 (reflecting a C->U editing event at the mRNA level). This percent of edited reads is shown in Figure 2e. Expanding beyond the single *DDOST* editing hotspot, we created the *ApoTrack* panel of RNA editing hotspots, which encompasses the top ~2000 most A3A-optimal RNA hairpins in the human transcriptome, and calculated in each sample the number of reads carrying evidence of a C->U editing event at one of these hotspot sites, divided by the total number of reads (in millions) covering these sites. This metric of edited reads per million is shown in Figure 2f.

### Quantitative RT-PCR assay for gene expression

Cells were seeded 24 hours before give a confluency of 80%. Cells were treated with drugs for 24 hours and RNA was extracted using the RNeasy Kit (Qiagen). cDNA was prepared from 500 ng total RNA with the First Strand Synthesis Kit (Invitrogen) using oligo-dT primers. Quantitative PCR was performed using FastStart SYBR Green (Roche) on a Lightcycler 480. mRNA expression relative to TBP mRNA levels was calculated using the delta-delta threshold cycle (ΔΔCT) method. Primer sequences were listed in Extended Data Table 3^57^.

### Droplet digital PCR assay

Purified RNAs were reverse transcribed using a High Capacity cDNA Reverse Transcription Kit (Thermo Fisher Scientific). 20 ng of cDNA, and indicated primers (2 μL) were added in the PCR reactions (ddPCR Supermix for Probes (No dUTP) mix from Bio-Rad) in a total of 25 μL. Then, 20 μL of the reaction mix was added to a DG8 cartridge (Bio-Rad), together with 70 μL Droplet Generation Oil for Probes (Bio-Rad) following by the generation of droplets using a QX200 Droplet Generator (Bio-Rad). Droplets were next transferred to a 96-well plate before to start the PCR reaction in thermal cycler (C1000 Touch Thermal Cycler, Bio-Rad) under the following conditions: 5 min at 95 °C, 40 cycles of 94 °C for 30 s, 53 °C for 1 min and then 98 °C for 10 min (ramp rate: 2 °C s−1). Droplets were analyzed with the QX200 Droplet Reader (Bio-Rad) for fluorescent measurement of fluorescein amidite (FAM) and hexachloro-fluorescein (HEX) probes. Gating was performed based on positive and negative DNA oligonucleotide controls. The ddPCR data were analyzed with QuantaSoft analysis software (Bio-Rad) to obtain fractional abundances of edited RNAs. Three or more biological replicates were analyzed for each sample. DDOST primers can be purchased from Bio-Rad (DDOST 558C #10031279, DDOST 558T #10031276).

### ATAC-seq

Untreated and gefitinib-treated PC9 cells were trypsinized and cryopreserved. ATAC-seq was performed by Active Motif. 100,000 nuclei were tagmented as previously described^58^, with some modifications based on Corces et al.^59^ using the enzyme and buffer provided in the Nextera Library Prep Kit (Illumina). Tagmented DNA was then purified using the MinElute PCR purification kit (Qiagen), amplified with 10 cycles of PCR, and purified using Agencourt AMPure SPRI beads (Beckman Coulter). Resulting material was quantified using the KAPA Library Quantification Kit for Illumina platforms (KAPA Biosystems), and sequenced with PE42 sequencing on the NextSeq 500 sequencer (Illumina). ATAC-seq reads in fastq format were trimmed to remove adapter sequences using skewer v0.2.2. Trimmed reads were aligned to hg19 using bowtie2 v2.3.3.1^60^ with the following parameters:’−D 20 −R 3 −N 1 −L 20 –local −i S,1,0.50 –rdg 5,1 –rfg 5,1 −X 2000’. Alignments were filtered using samtools v1.9^61^ to remove duplicate reads and retain only proper pairs with mapping quality greater than 30. Reads mapping to blacklisted regions were removed using samtools v1.9. Reads were converted to bed format and reads on the forward and reverse strands were shifted +4 and −5 bp respectively. MACS2 v2.1.1^62^ was then used to call peaks and summits using the following parameters:’–nomodel –extsize 200 –shift −100 –nolambda –keep-dup all −q 0.1’. IDR v2.0.4.2 was used to filter enriched regions at an IDR of less than 0.05 to produce high-confidence peak and summit sets. High-confidence summits were expanded to 500bp and a count matrix was generated with Rsubread v1.30.9^63^. This matrix was imported to R and chromVAR^64^ was used in conjunction with motifs in the JASPAR2018 (10.18129/B9.bioc.JASPAR2018)^65^ and chromVARmotifs (https://github.com/GreenleafLab/chromVARmotifs) R packages to generate motif-level measures of differential accessibility. Signal tracks were generated from bigwig files created from aligned bams using Deeptools v3.1.1 bamCoverage command^66^ with the following parameters: “–Offset 1 −bs 50 –smoothLength 150 –maxFragmentLength 2000 –scaleFactor SCALEFACTOR”. Scaling factors for each sample were generated by quantifying total cutsites within each sample’s IDR-filtered peaks and dividing by 1e6.

### Comet assay

single-cell gel electrophoresis was carried out by using Trevigen’s comet assay kit (#4250-050−K) with slight modifications. Briefly, PC9 cell suspension (4 × 10^5^/mL) and melted LM agarose (at 37°C) were prepared in 1:10 (v/v) ratio. 50 μL of this solution was poured onto the comet slide. Slides were then kept at 4°C in the dark for 10 min and afterwards at 37°C for 5 min for better agarose adhesion. The slides were placed in an ice-cold lysing solution for 90 min at 4°C. After lysis, slides were immersed in freshly prepared neutral electrophoresis buffer (pH =9, 500 mM Tris base, 150 mM Sodium acetate) for 30 min at 4°C in the dark. Subsequently, the slides were placed in an electrophoresis chamber filled with cold neutral buffer and run for 60 min at 15V in cold room. After the run, slides were incubated in DNA precipitation solution (Ammonium acetate 1 M in 95% ethanol) for 30 min at room temperature and then incubated in 70% ethanol for 30 min at room temperature. Slides were dried at 37°C until agarose completely disappeared. Comets were stained with SYBR™ Gold (Invitrogen #S7563) for 30 min at room temperature. Then slides were rinsed in water and completely dried at 37°C. Finally, slides were mounted with SlowFade™ Diamond Antifade Mountant (Invitrogen #S36963). Slides were imaged with Echo revolve microscope and pictures were analyzed with Open Comet ImageJ plug-in^67^. Quantification of the comet tail moment was calculated by multiplying the % of DNA in the tail by the distance between means of the head and tail distributions.

### Immunofluorescence based cell-cycle profile

PC9 cells were grown on coverslips. Before performing the staining, cells were incubated in media containing 5-ethynyl-2’-deoxyuridine 10 μM for 60 min at 37°C. Cells were then washed twice with cold PBS and fixed 15 min at room temperature with PBS 3.7% formaldehyde. After fixation, cells were permeabilized with PBS 0.5% Triton−X 100 for 10 min at room temperature. Afterwards cells were washed twice with PBS 0.1% Tween and blocked for 60 min at room temperature with blocking solution (PBS, 2.5% bovine serum albumin, 0.1% Tween). Both primary and secondary antibodies were diluted in blocking solution. After 3 hours incubation with primary antibody at room temperature, cells were washed three times with PBS 0.1% Tween and incubated with secondary antibody for 60 min at room temperature. Finally, EdU detection was performed by using Click-iT™ EdU Alexa Fluor™ 647 Imaging Kit (Thermo scientific #C10340) following manufacturer’s instruction. Coverslips were mounted with SlowFade™ Diamond Antifade Mountant with Dapi (Invitrogen #S36964) and imaged with Echo revolve microscope. Images were analyzed and fluorescence was quantified with MATLAB software. In brief, cell nuclei were segmented using a custom-made image processing pipeline that is able to distinguish them from the background. The pipeline identifies the nuclei based on Dapi staining, their size and circularity. Nuclei that are close to image borders or that are too close and cannot be individually segmented were automatically removed by the software. Both γH2AX and Edu fluorescence intensities were quantified by the software only within the segmented nuclei.

### Doxycycline inducible APOBEC3A overexpression

cDNA was synthesized by GenScript with a beta-globin intron between exon 2 and 3 of APOBEC3A and a Flag tag at C-terminus. The plasmid expressing Flag-APOBEC3A was generated by inserting the cDNA into the pInducer20 vector using the Gateway Cloning System (Thermo Fisher Scientific)^68^. Cells were seeded into six-well plates at a density of 2 × 10^5^ cells per well. 24 hours later, cells were infected with APOBEC3A wild-type or catalytically inactive mutant APOBEC3A^E72A^ viral particles for 24 hours at 37°C with 8 μg/mL of polybrene (Millipore #TR-1003-G). PC9 cells were selected in 600 μg /mL of G418 (Gibco #10131035) for 4 days. Cells were incubated with 200 ng/mL of doxycycline (Sigma) for 72-96 hours. The expression of APOBEC3A was confirmed by western blotting as previously described^69^ (Extended Data Figure 3b).

### APOBEC3A lentiviral shRNA knockdown

Non-targeting scrambled (SCR) control shRNA and APOBEC3A specific shRNA (Hairpin sequence: 5’-CAGTACCAGACTCCATCTCAA-3’) were delivered on the pLKO.1-background vector (from MGH Molecular Profiling Laboratory) and packaged using 293T cells. PC9 cells were seeded into six-well plates at a density of 2 × 10^5^ cells per well. 24 hours later, infected with viral particles for 24 hours at 37°C with 8 μg/mL of polybrene. PC9 cells were selected in 2 μg/mL of puromycin for 4 days. APOBEC3A knockdown efficiency was confirmed by quantitative RT-PCR.

### CRISPR/Cas9 knockout of APOBEC3A

Plasmid pX458 (Addgene #48138) was kindly donated by Feng Zhang^70^ and used for expression of the gRNA and the human codon-optimized SpCas9 protein. Target sequences for CRISPR interference were designed using the sgRNA designer developed by BROAD institute (http://portals.broadinstitute.org/gpp/public/analysis-tools/sgrna-design). A non-targeting sgRNA from the Gecko library v2 was used as a scramble sgRNA. sgRNA target sequences are used: APOBEC3A 5’-GACCTACCTGTGCTACGAAG-3’; scramble 5’-ATCGTTTCCGCTTAACGGCG-3’. Complementary oligonucleotides encoding the gRNAs targeting APOBEC3A sequence were annealed and ligated into pX458. One day prior to transduction, 1 × 10^5^ cells were seeded into a single well of a 24 well plate. The cells were transfected with Lipofectamine 3000 Reagent (Thermo Fisher Scientific) according to the manufacturer’s protocol. Four days after transduction, GFP-positive cells were sorted by flow cytometry. The knockout efficacy was confirmed by direct sequencing.

### Crystal Violet

Cells were seeded to give a confluency of ~80%. Cells were drugged the following day and subsequently media and drug were replaced twice per week. After 4-6 weeks in drug, cells were fixed with glutaraldehyde and stained with 0.1% crystal violet.

### RealTime-Glo viability assay

Cell viability was assayed in situ once a week starting the day after seeding, using the RealTime-Glo assay (Promega) according to the manufacturer’s protocol. Briefly, MT Cell Viability Substrate and NanoLuc Enzyme were diluted 1:500 in medium, and 25 μl was added to each well (1/5 total final volume). Cells were incubated for 1 hour at 37 °C and luminescence was measured. Fresh medium containing drug was used to replace the assay reagents immediately after each assay.

### Mouse xenograft studies

All mouse studies were conducted through Institutional Animal Care and Use Committee– approved animal protocols in accordance with institutional guidelines (MGH Subcommittee on Research Animal Care, OLAW Assurance A3596-01). For xenograft studies, cell line suspensions were prepared in 1:10 matrigel and 5 × 10^6^ cells were injected subcutaneously into the flanks of female athymic nude (Nu/Nu) mice (6-8 weeks old). Visible tumors developed in approximately 2-4 weeks. Tumors were measured with electronic calipers and the tumor volume was calculated according to the formula Vol = 0.52 × L × W^2^. Mice with established tumors were randomized to drug treatment groups using covariate-adaptive randomization to minimize differences in baseline tumor volumes: osimertinib at 5 mg/kg (10% 1-methyl-2-pyrolidone, 90% PEG300). Drug treatments were administered by oral gavage and tumor volumes were measured twice weekly, as described above. Mice were treated until tumors had reached stable minimal residual disease state defined by no change in tumor volume for at least 2 consecutive measurements. Tumors for RNA analyses were snap-frozen in liquid nitrogen immediately upon harvesting. Investigators performing tumor measurements were not blinded to treatment groups.

### Data and statistical analysis

Data were analyzed using GraphPad Prism software (GraphPad Software). Unless otherwise specified, data displayed are mean ± s.e.m. Pairwise comparisons between groups (for example, experimental versus control) were made using paired or unpaired Student’s t-tests as appropriate.

### Data availability

Data availability. All data presented in this manuscript are available from the corresponding author upon reasonable request. Sequencing has been deposited at the GEO (http://www.ncbi.nlm.nih.gov/geo) under the accession number GSE75602 and GSE114647.

## Acknowledgements

We thank all members of the Hata and Benes Lab for helpful discussions and feedback. This study was funded by support from the NIH K08 CA197389 (A.N.H), R01 CA249291 (A.N.H), R01 CA137008 (L.V.S.), U01CA220323 (N.J.D.), Doris Duke Charitable Foundation Clinical Scientist Development Award (A.N.H), Smith Family Foundation Award (A.N.H), SU2C/NSF/V Foundation Convergence Award (A.N.H and L.V.S), the Ludwig Center at Harvard (A.N.H.), Tosteson & FMD Award (H.I.), the Lungstrong foundation, Targeting a Cure for Lung Cancer, and Be a Piece of the Solution, the Landry Family and the Suzanne E. Coyne Family.

## Author Contributions

H.I., A.N.H, M.S.L. designed the study, analyzed the data and wrote the paper. H.I., N.N., W.S., M.S., H.F.C., F.M.S., P.J., S.O., D.T., H.A., V.N. and A.N.H performed cell line and biochemical studies. N.P., S.B., M.G.C. performed tumor xenograft studies. K.D., A.R. generated the patient derived cell lines. A.A., A.L., M.L., C.O., C.S.C., J.J.L., Y.M., and M.S.L performed computational analysis. L.Z., N.J.D., C.B., G.G., R.B., J.A.E. were involved with study design. M.K.B., R.G.C., A.T.S., J.J.L, L.V.S. and Z.P. provided NSCLC patient samples. A.N.H and M.S.L. contributed equally to the study. All authors discussed the results and commented on the manuscript.

---

**Extended Data Figure 1.**
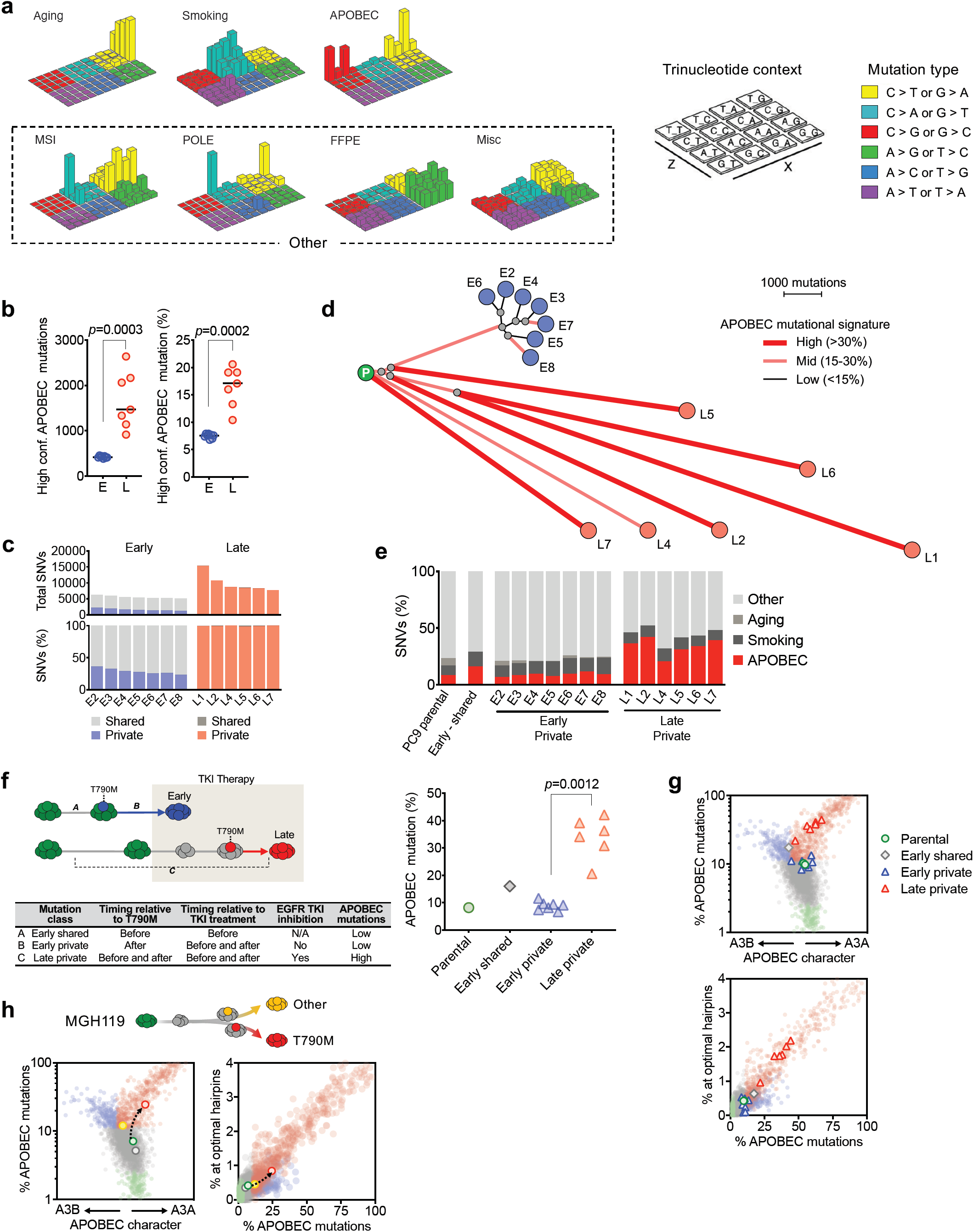
TKI-resistant clones that evolve from drug-tolerant persister cells accumulate APOBEC mutations. **a**, Lego plots of the 7 mutational signatures resolved by NMF with assigned biological process. For simplicity, MSI, POLE, FFPE and Miscellaneous signatures are combined and plotted as “Other” throughout the manuscript. **b**, Number and percentage of <u>T</u>CT→<u>T</u>GT and <u>T</u>CA→<u>T</u>GA mutations that are highly specific for APOBEC. **c**, Shared and private mutations in early and late resistant clones. **d**, Phylogenetic tree depicting evolutionary relationships of early and late resistant clones based on pattern of shared and private mutations. **e**, Mutational signatures of private and shared mutations. **f**, Relationship between timing of T790M/TKI treatment and APOBEC mutational signatures for shared and private mutations. **g**, A3A character and hairpin mutation frequency of shared and private mutations. PCAWG reference samples are colored as in Figure 1g. **h**, A3A character and hairpin mutation frequency of MGH119 mutations.

**Extended Data Figure 2.**
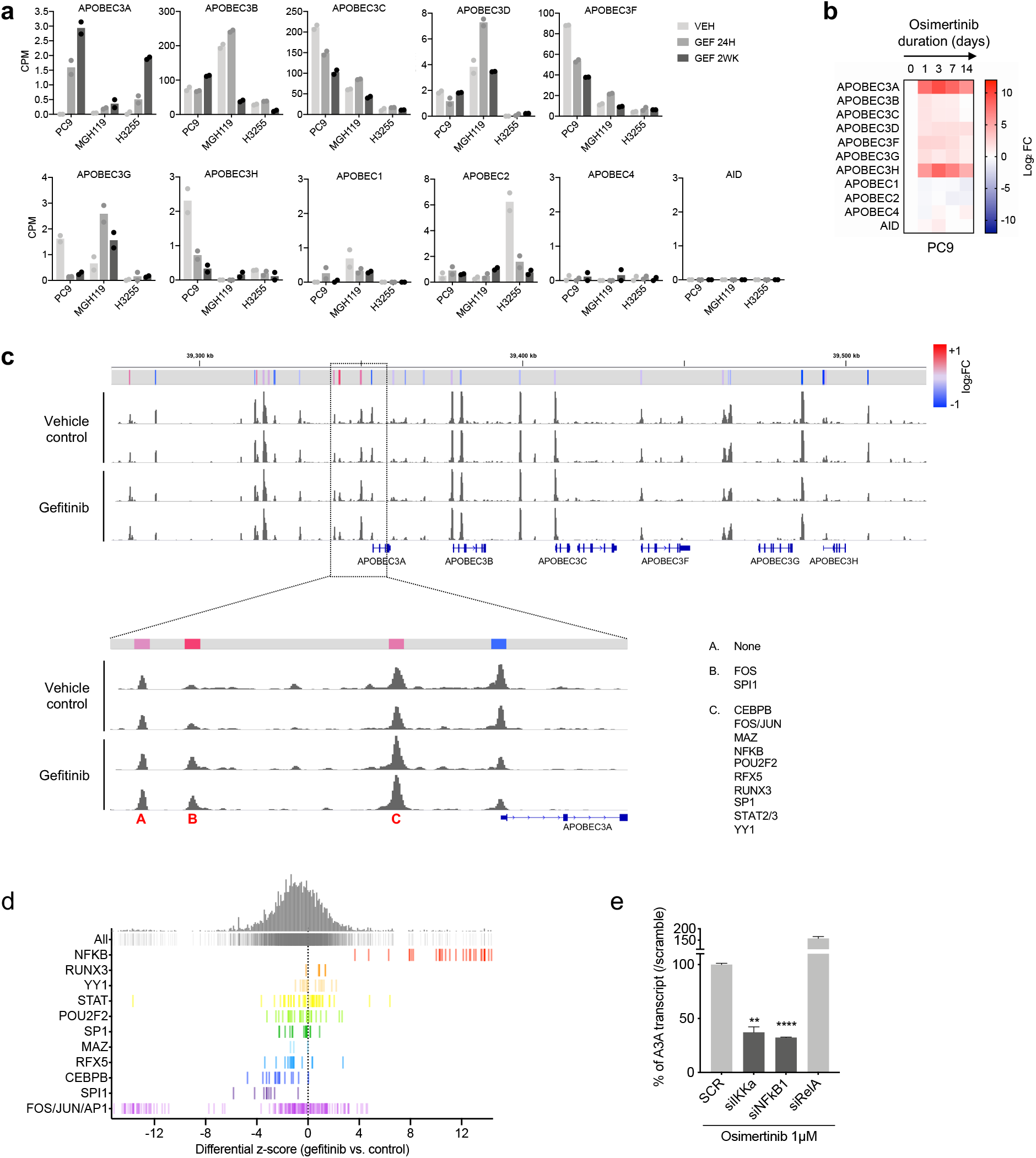
Lung cancer targeted therapies induce APOBEC3A expression. **a**, RNA expression levels of APOBEC family genes determined by RNA-seq in EGFR NSCLC cell lines treated with 300 nM gefitinib for 0, 1 and 14 days. Data are from GSE114647^71^. **b**, Expression of APOBEC family genes in response to osimertinib. Cells were treated with 1 μM osimertinib for up to 14 days and gene expression was determined by quantitative RT-PCR. Data are expressed as log_2_ fold change relative to untreated control. **c**, Chromatin accessibility profile of APOBEC3 locus (chromosome 22) of PC9 cells treated with 300 nM gefitinib for 0 or 14 days, with expanded view of A3A promoter region. Relative fold-change of peak accessibility in treated vs un-treated cells is shown. Transcription factors mapping to these peak regions were identified from the ENCODE database. **d**, Differential z-score (gefitinib vs control) of global motif accessibility scores for each identified transcription factor family. **e**, Osimertinib-induced APOBEC3A transcript level after siRNA knockdown of indicated genes. Data are expressed as a percentage of APOBEC3A transcript level in control cells (SCR) transfected with a non-targeting siRNA after 24 hours osimertinib treatment.

**Extended Data Figure 3.**
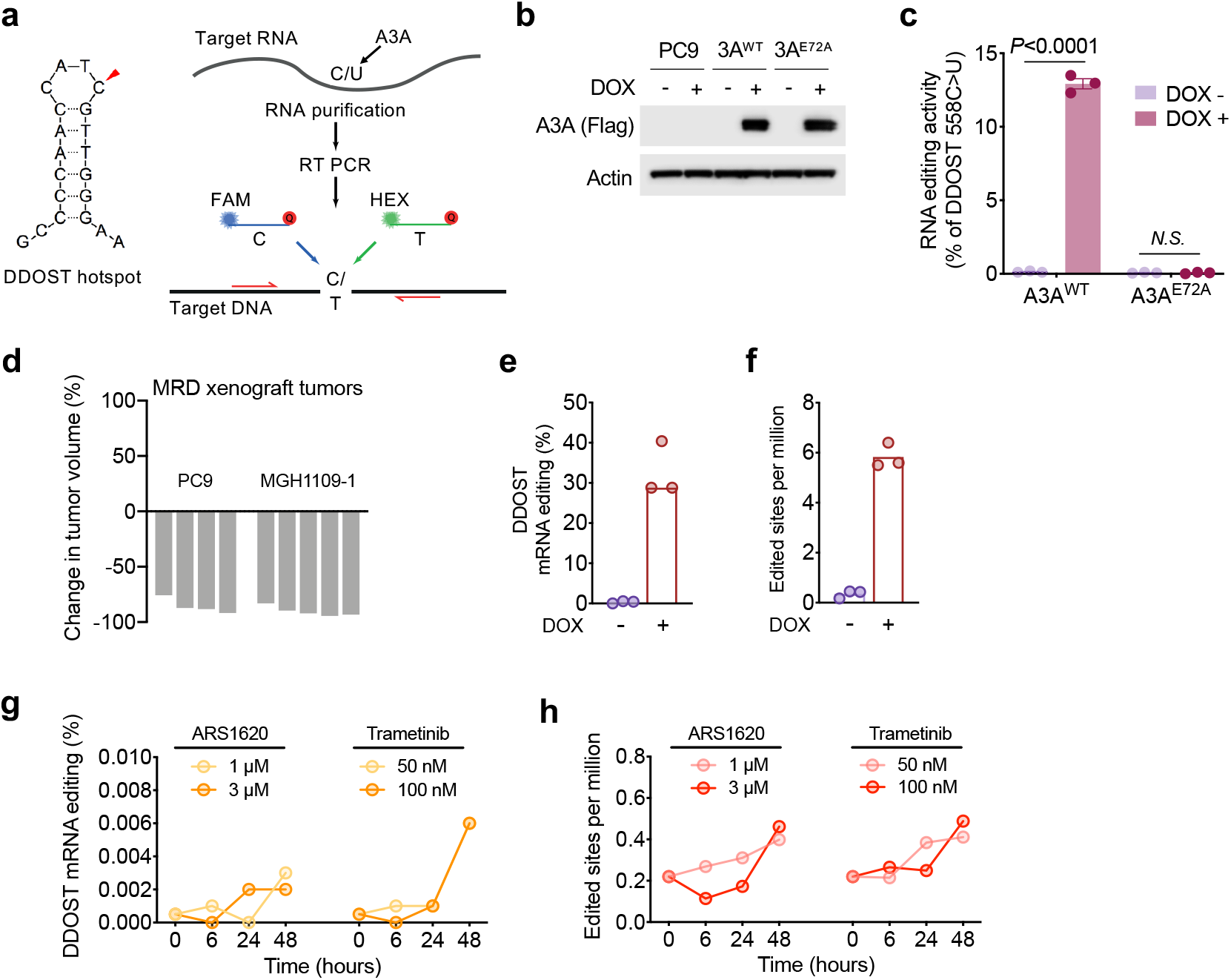
TKI treatment induces APOBEC3A RNA editing. **a**, Schema for allele specific droplet digital PCR assay for quantitating A3A editing at the DDOST hairpin hotspot (adapted from Jalili et al.^27^). **b-c**, Overexpression of wild-type A3A but not the catalytic inactive A3A^E72A^ mutant induces A3A editing of DDOST. PC9 cells transduced with doxycycline-inducible wild-type A3A or the catalytic inactive mutant A3A^E72A^ were treated with doxycycline to induce expression of A3A and DDOST editing was determined by droplet digital PCR (mean ± s.d., 3 biological replicates). **d**, Change in volume of osimertinib treated EGFR mutant NSCLC xenograft tumors at the time of harvesting minimal residual disease (MRD) for DDOST hairpin hotspot analysis in Figure 2d. **e**, PC9 cells transduced with dox-inducible wild-type A3A were treated with or without doxycycline and subjected to mRNA-seq. % of DDOST editing was determined by dividing the number of edited RNA-seq reads by the total number of reads aligning to the DDOST hotspot. **f**, ApoTrack was applied to RNA-seq data from PC9 cells after doxycycline-induction of wild-type A3A (see Extended Data Figure 3b,c) to quantify reads containing UCN > UUN mutations in hairpin loops of sequence NUC or NNUC at the end of stably-paired stems at ~2000 sites across the transcriptome. A3A editing is expressed as the number of edited reads per 1000 reads for all genes combined. **g-h**, mRNA-seq was performed on H358 *KRAS^G12C^* NSCLC cells treated with ARS-1620 or trametinib to quantify DDOST and global transcriptome editing (ApoTrack).

**Extended Data Figure 4.**
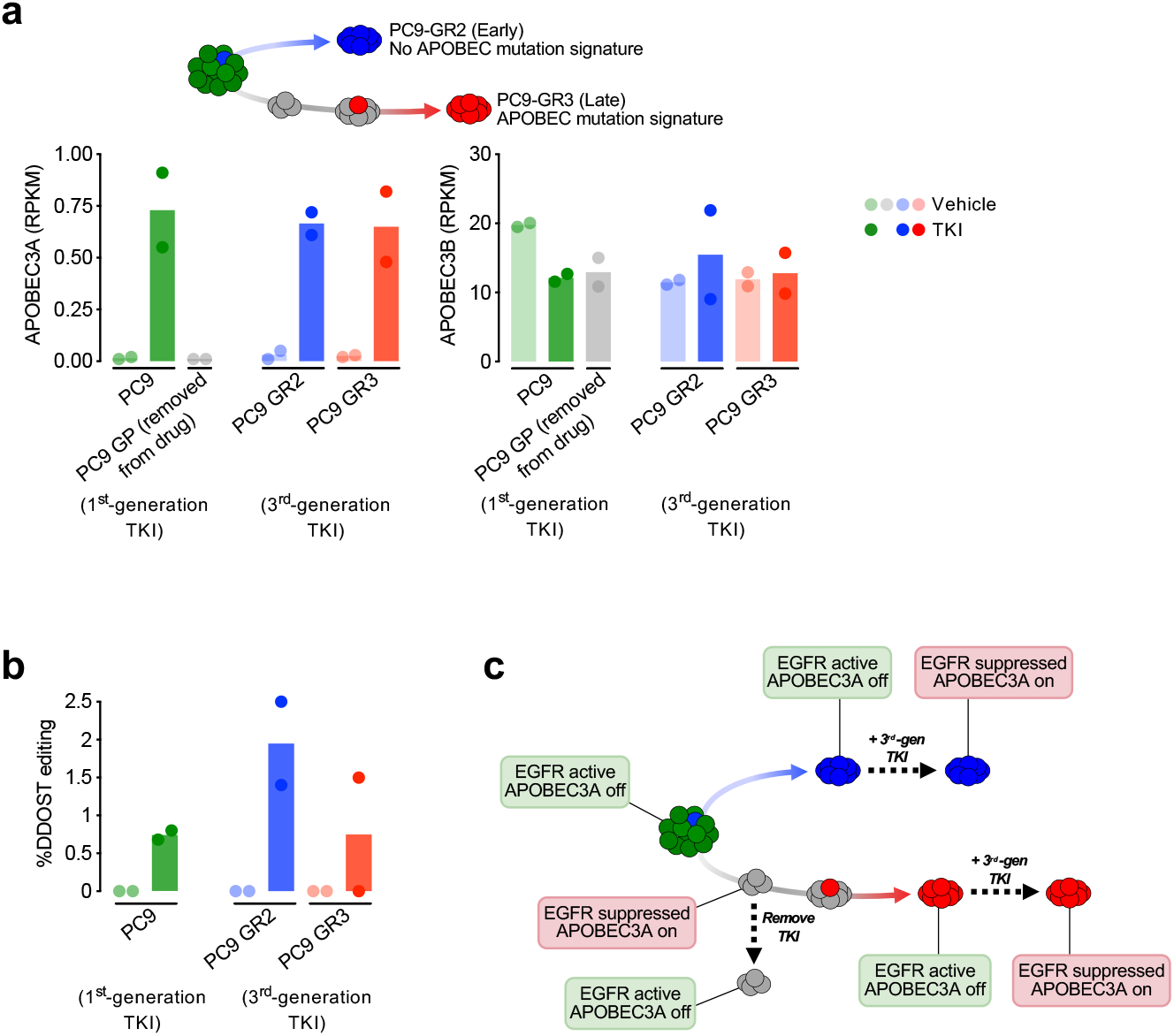
APOBEC editing coincides with suppression of EGFR signaling during evolution of acquired resistance. **a**, A3A and A3B expression levels from RNA-seq (GSE75602) performed on parental PC9 cells, PC9 drug-tolerant persister cells after 2 weeks of gefitinib treatment (GP), early *EGFR^T790M^* resistant clone PC9-GR2 and late *EGFR^T790M^* resistant clone PC9-GR3 (previously described in Hata and Niederst et al. ^14^). PC9 cells were treated with gefitinib, PC9-GR2/GR3 cells were treated with the third generation EGFR inhibitor WZ4002 (all for 24 hours). **b**, Percentage of DDOST hotspot reads with A3A editing in PC9 and PC9-GR2/GR3 cells treated with gefitinib or WZ4002, respectively. **c**, Summary of the relationship between EGFR activity and A3A expression/activity during the evolutionary trajectories of PC9 acquired resistance.

**Extended Data Figure 5.**
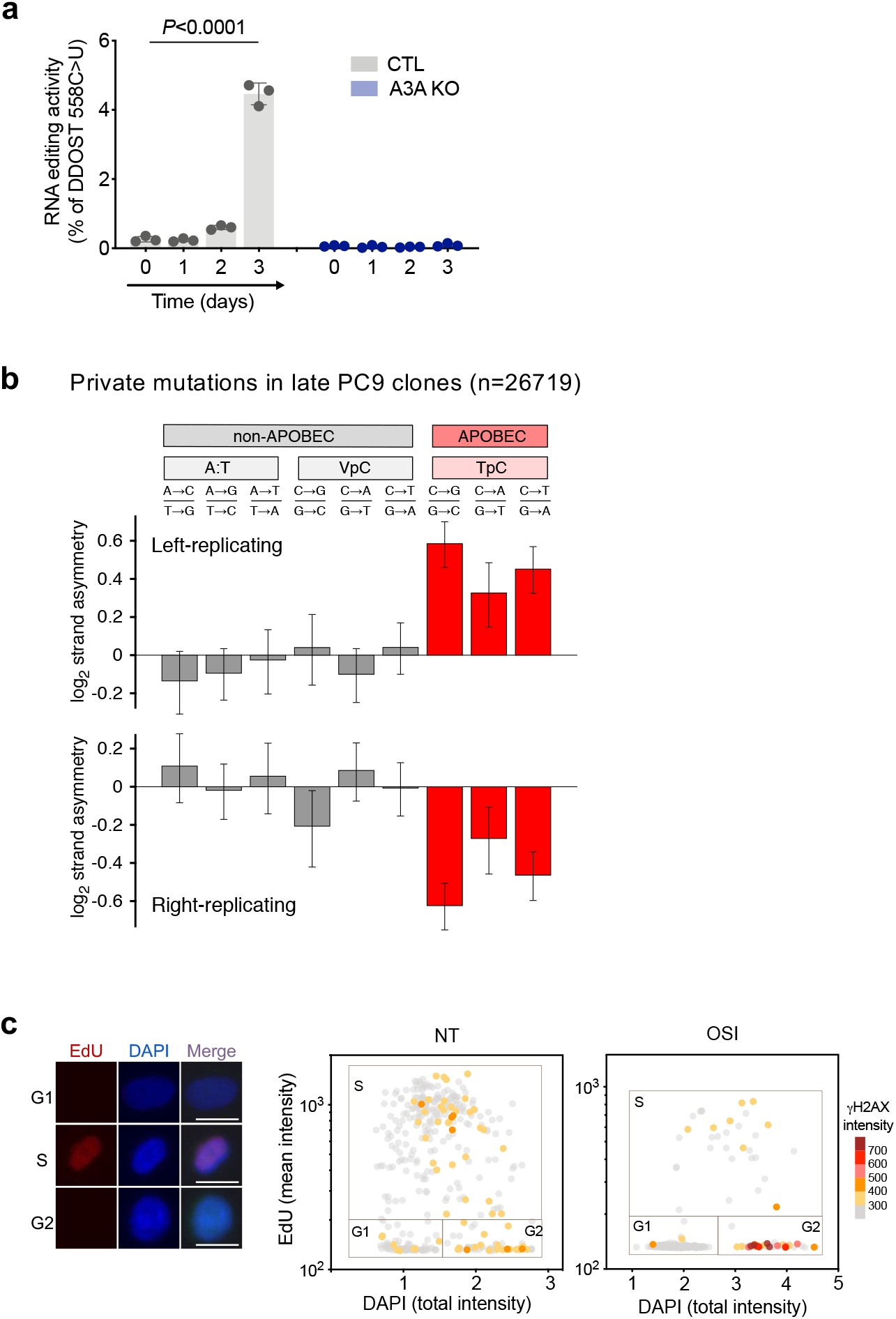
TKI-induced APOBEC3A leads to genomic instability and facilitates evolution of drug-resistant clones. **a**, PC9 A3A knockout (KO) cells exhibit no TKI-induced DDOST RNA editing. PC9 control and PC9 A3A KO cells were treated with or without 1 μM osimertinib for up to 3 days and DDOST editing was determined by droplet digital PCR. (mean ± s.d. 3 biological replicates) **b**, A3A mutational signatures in late evolving PC9 resistant clones exhibit replication asymmetry. **c**, Representative G1, S and G2 cells; scale bars = 10 μm (Left panel). Scatter plots of EdU cell cycle assay in Figure 3e (Right panel). PC9 cells were treated with 1 μM osimertinib for 14 days and stained with EdU/DAPI to resolve cell cycle phase, and γH2AX to quantify DNA damage (NT = no treatment).

**Extended Data Figure 6.**
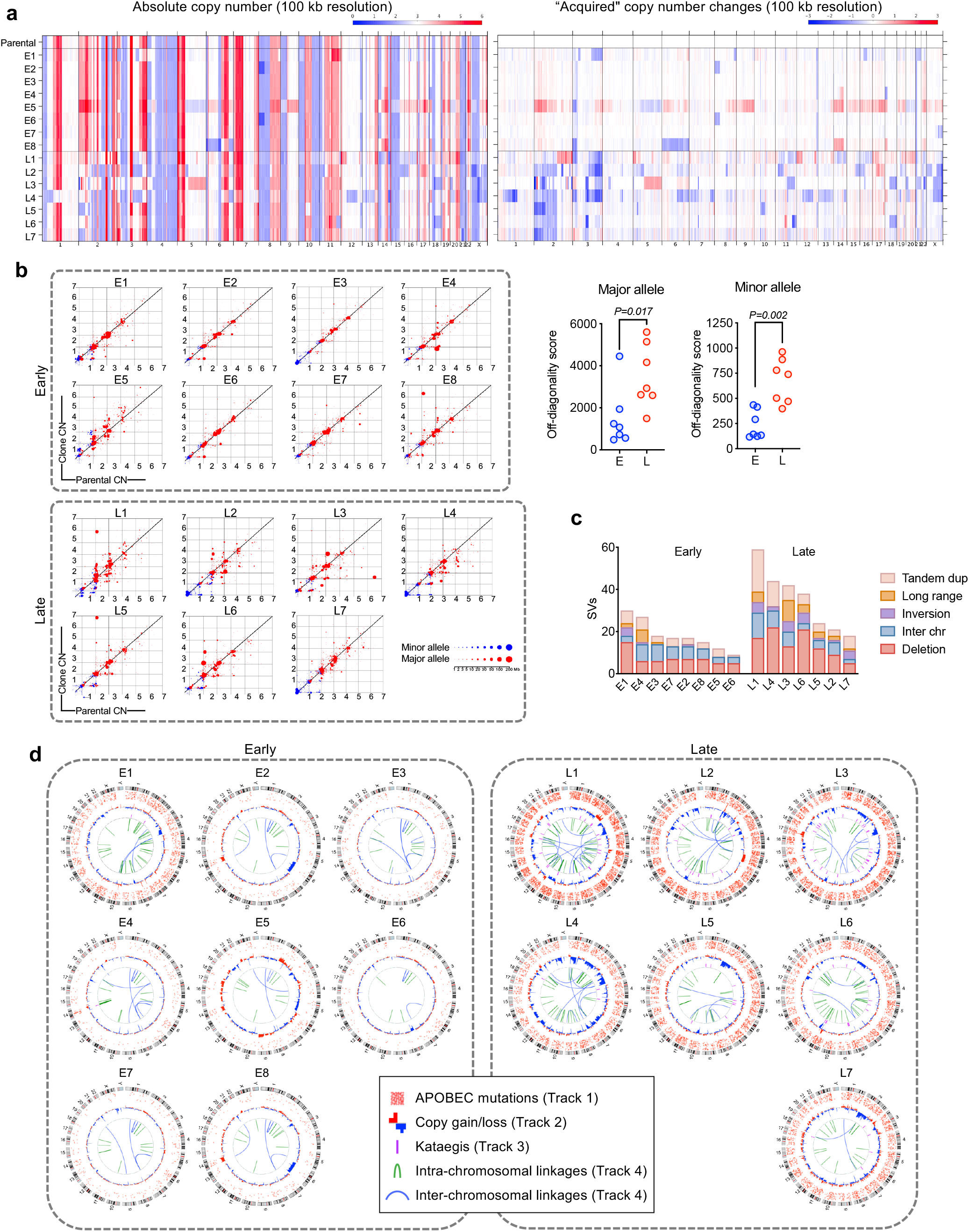
Late-evolving PC9 resistant clones have evidence of increased genomic instability. **a**, Copy number profiles of early and late resistant PC9 clones. **b**, Acquired copy number changes in each sample are shown on scatter plots where the y-axis is the copy number in the sample, and the x-axis is the copy number in the parental sample it was derived from. Genomic copy number segments are shown for both the minor allele (blue) and major allele (red). Size of points corresponds to the size of the copy-number segments. Points along the diagonal represent genomic segments that did not change in copy number between the parental and derived sample. Points above and below the diagonal represent copy-number gains and losses, respectively, relative to the parental sample. **c**, Late evolving PC9 clones have increased structural variations (SVs) compared with early resistant clones. SVs were determined relative to parental cells using dRanger and Breakpointer. **d**, Circos plots depicting APOBEC mutations, copy number changes, kataegis events and intra/inter-chromosomal interactions in PC9 resistant clones (all relative to parental cells).

**Extended Data Figure 7.**
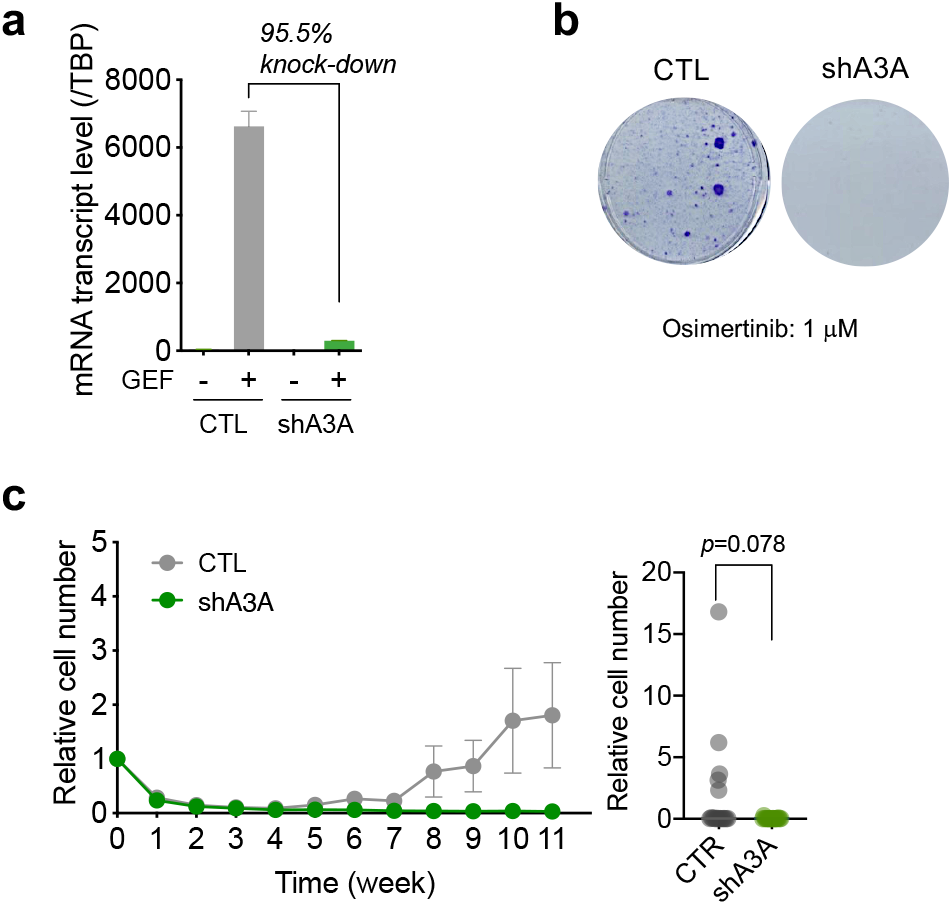
Knockdown of A3A by shRNA suppresses the emergence of drug-tolerant and resistant clones. **a**, Knockdown efficacy of A3A shRNA. PC9 shA3A cells have > 95% reduction in TKI-induced A3A gene expression compared with control shRNA cells. Cells were treated with 1 μM gefitinib for 24 hours. **b**, Crystal violet staining of PC9 shA3A and control cells treated with 1 μM osimertinib for 4 weeks. **c**, Long-term monitoring of emergence of resistant clones during osimertinib treatment. PC9 shA3A and control cells were treated with 1 μM osimertinib for 11 weeks and cell number was assessed using RealTime-Go (N=18 independent pools each, mean ± s.e.m.). Right panel shows the cell number of each pool at week 11.

**Extended Data Figure 8.**
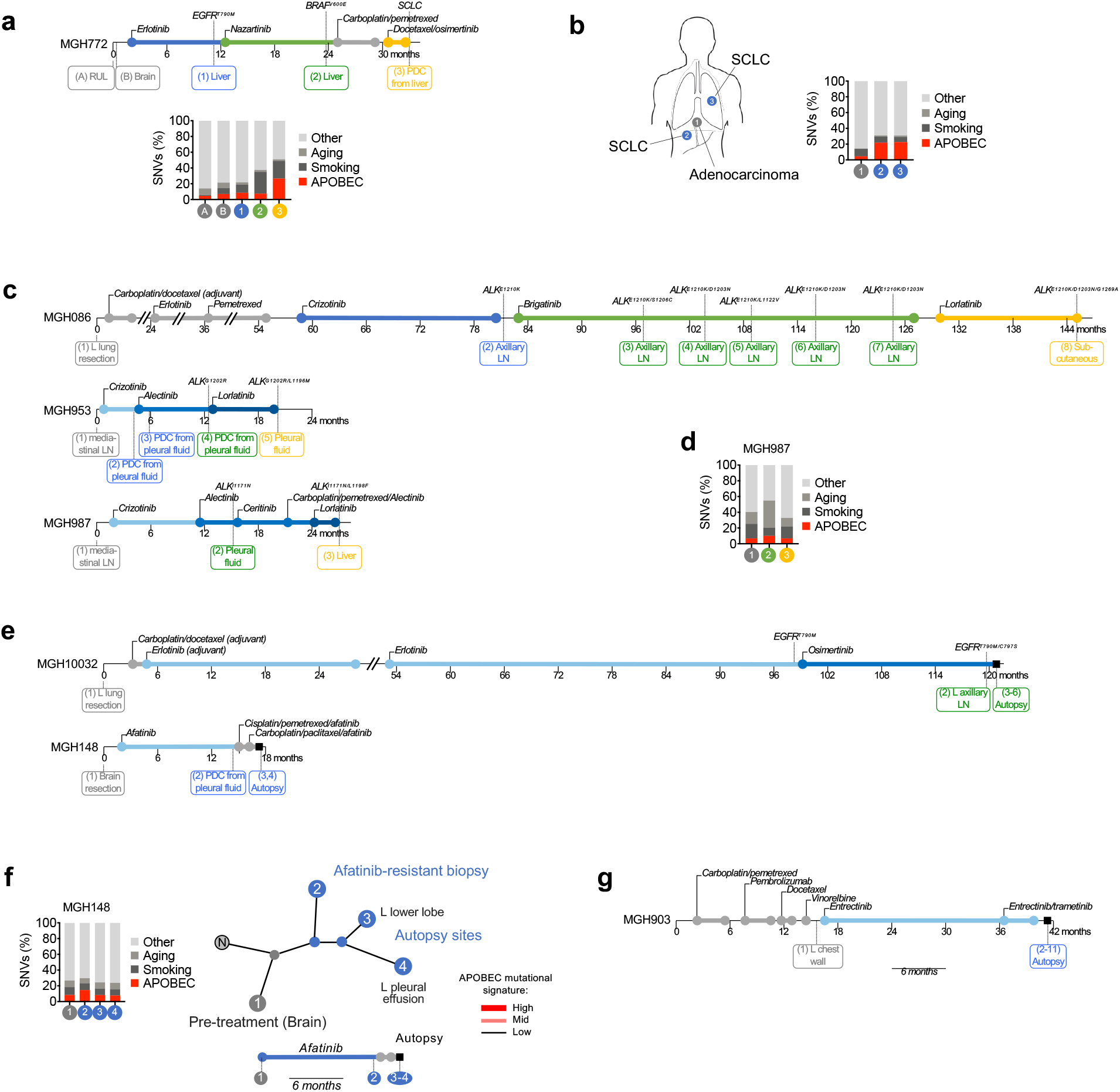
APOBEC3A mutational signatures in NSCLC patients with acquired TKI resistance. **a**, Clinical history, mutational signatures and clonal relationship of serial biopsies from patient MGH772^43^. **b**, Clinical history, mutational signatures and clonal relationship of metastatic sites from MGH7 autopsy^42^. **c**, Clinical histories of ALK NSCLC patients who acquired compound ALK resistance mutations after treatment with sequential ALK TKIs^17^. **d**, Mutational signatures in MGH987 sequential biopsies. **e**, Clinical histories of MGH10032 and MGH148 EGFR NSCLC autopsy cases. **f**, Mutational signatures and clonal relationship of metastatic sites from MGH148. **g**, Clinical history of MGH903 NTRK NSCLC autopsy case.

**Extended Data Figure 9.**
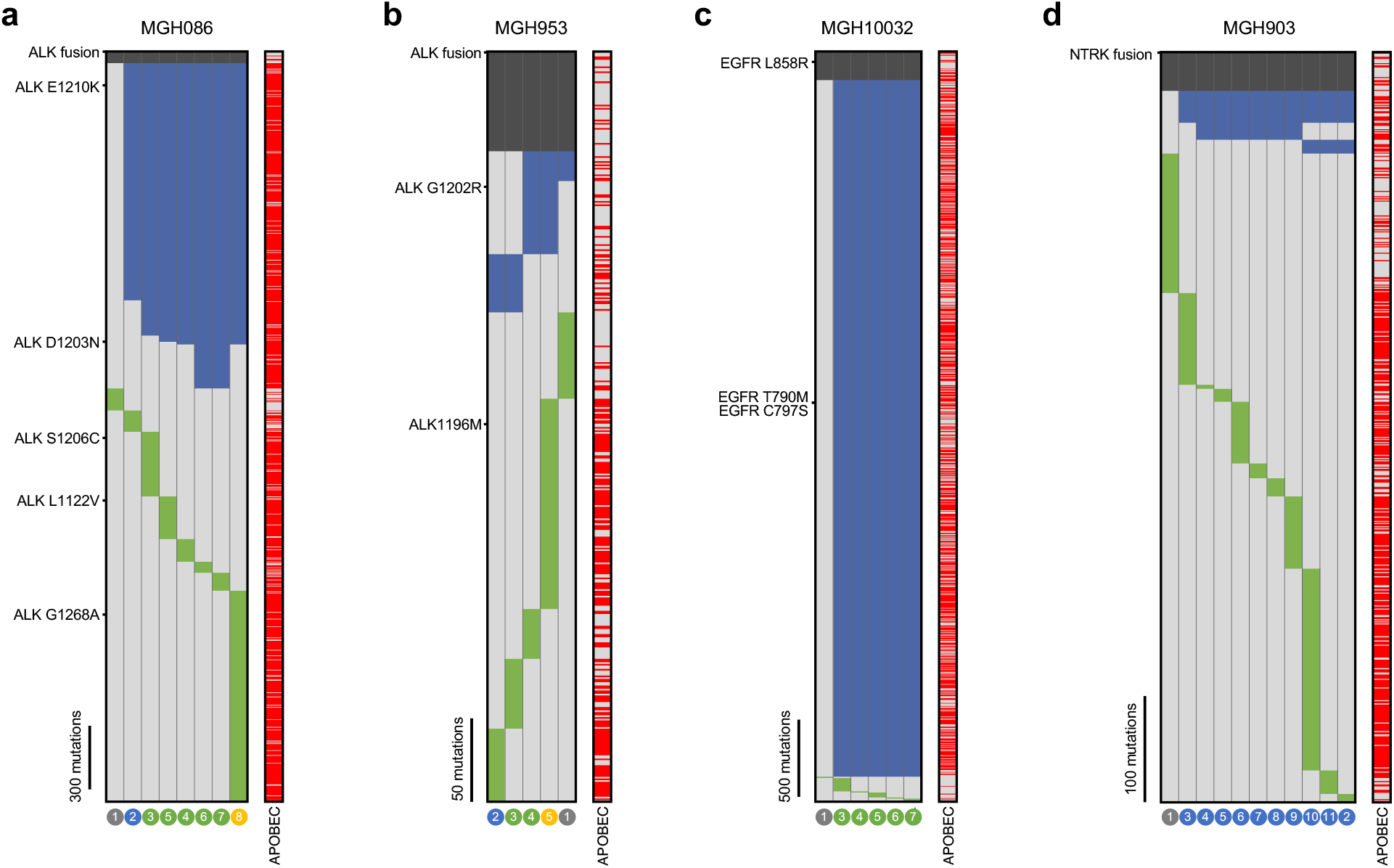
Mutational heterogeneity in sequential biopsies and autopsy samples. Heatmaps depict presence of mutations in each sample from MGH086 (**a)**, MGH953 **(b)**, MGH10032 **(c)**and MGH903 **(d)**. Samples are denoted as in Figure 4 and Extended Data Figure 8. Truncal mutations present in all samples are colored dark gray, shared mutations common to 2 or more samples are colored blue, and private mutations are colored green. APOBEC mutations are indicated in red. The absence of a mutation is denoted by light gray.

