## Supplemental Tables for "APOBEC3A drives acquired resistance to targeted therapies in non-small cell lung cancer"

Extended Data Table 1. Cell line characteristics.

| Cell line | Oncogene | Targeted Rx | Resistance mechanism |
|---|---|---|---|
| PC9 | EGFR exon 19 del | - | Naïve |
| H3255 | EGFR exon 21, L858R | - | Naïve |
| MGH119-1 | EGFR exon 19 del | - | Naïve |
| MGH121-1 | EGFR exon 19 del | Erlotinib | EGFR T790M |
| MGH134-1 | EGFR exon 21 L858R | Erlotinib | EGFR T790M |
| MGH141-1 | EGFR exon 19 del | Erlotinib | EGFR T790M |
| MGH792-1 | EGFR exon 21 L858R | Erlotinib | On-treatment |
| MGH840-1 | EGFR exon 19 del | - | Naïve |
| MGH1074-1 | EGFR exon 19 del | - | Naïve |
| MGH1109-1 | EGFR exon 19 del | - | Naïve |
| MGH1144-2 | EGFR exon 19 del | - | Naïve |
| MGH1157-1 | EGFR exon 19 del | - | Naïve |
| MGH10026-1 | EGFR exon 21 L858R | - | Naïve |
| H3122 | EML4-ALKv1 | - | Naïve |
| H2228 | EML4-ALKv3 | - | Naïve |
| MGH006-1 | EML4-ALKv1 | - | Naïve |
| MGH026-1 | EML4-ALKv3 | - | Naïve |
| MGH064-1 | EML4-ALKv2 | - | Naïve |
| H358 | KRAS G12C | - | Naïve |

Extended Data Table 2. Mutation number and kataegis events in reisitant models.

| Cell line | | Hata et al. Nature Med. 2016 | APOBEC | Smoking | Aging | Other | Kataegis |
|---|---|---|---|---|---|---|---|
| Early T790M PC9 clones | E1 | PC9-GR2 | 783 | 1172 | 1 | 4768 | 0 |
| | E2 | Early T-23 | 671 | 959 | 73 | 4604 | 0 |
| | E3 | Early T-24 | 680 | 919 | 43 | 4283 | 0 |
| | E4 | Early T-25 | 668 | 890 | 2 | 4024 | 10 |
| | E5 | Early T-26 | 621 | 895 | 6 | 3845 | 0 |
| | E6 | Early T-1 | 640 | 880 | 26 | 3691 | 0 |
| | E7 | Early T-12 | 662 | 826 | 9 | 3719 | 0 |
| | E8 | Early T-19 | 608 | 858 | 9 | 3609 | 0 |
| Late T790M PC9 clones | L1 | Late T-1 | 4938 | 2018 | 0 | 8468 | 16 |
| | L2 | Late T-4 | 4040 | 1458 | 2 | 5265 | 32 |
| | L3 | PC9-GR3 | 4142 | 1350 | 2 | 5014 | 19 |
| | L4 | Late T-2 | 1545 | 1271 | 3 | 5949 | 3 |
| | L5 | Late T-5 | 2286 | 1212 | 0 | 5076 | 8 |
| | L6 | Late T-9 | 2485 | 1001 | 1 | 4866 | 16 |
| | L7 | Late T-3 | 2677 | 930 | 2 | 4078 | 8 |

Extended Data Table 3. Quantitative PCR primer information.

| Gene Symbol | mRNA NCBI Accession | 5’ Primer Seq (5'-3') | 3’ Primer Seq (5'-3') |
|---|---|---|---|
| **APOBECs** | | | |
| APOBEC1 | NM_001644 | gggaccttgttaacagtggagt | ccaggtgggtagttgacaaaa |
| APOBEC2 | NM_006789 | aagtagggcaactgggcttt | ggctgtacatgtcattgctgtc |
| APOBEC3A | NM_145699 | gagaagggacaagcacatgg | tggatccatcaagtgtctgg |
| APOBEC3B | NM_004900 | gaccctttggtccttcgac | gcacagccccaggagaag |
| APOBEC3C | NM_014508 | agcgcttcagaaaagagtgg | aagtttcgttccgatcgttg |
| APOBEC3D | NM_152426 | acccaaacgtcagtcgaatc | cacatttctgcgtggttctc |
| APOBEC3F | NM_145298 | ccgtttggacgcaaagat | ccaggtgatctggaaacactt |
| APOBEC3G | NM_021822 | ccgaggacccgaaggttac | tccaacagtgctgaaattcg |
| APOBEC3H | NM_181773 | agctgtggccagaagcac | cggaatgtttcggctgtt |
| APOBEC4 | NM_203454 | ttctaacacctggaatgtgatcc | tttactgtcttctagctgcaaacc |
| AID | NM_020661 | gactttggttatcttcgcaataaga | aggtcccagtccgagatgta |
| **Reference Gene** | | | |
| TBP | NM_003194 | cccatgactcccatgacc | tttacaaccaagattcactgtgg |
